# Searching across-cohort relatives in 54,092 GWAS samples via encrypted genotype regression

**DOI:** 10.1101/2022.10.18.512407

**Authors:** Qi-Xin Zhang, Tianzi Liu, Xinxin Guo, Jianxin Zhen, Meng-yuan Yang, Saber Khederzadeh, Fang Zhou, Xiaotong Han, Qiwen Zheng, Peilin Jia, Xiaohu Ding, Mingguang He, Xin Zou, Jia-Kai Liao, Hongxin Zhang, Ji He, Xiaofeng Zhu, Daru Lu, Hongyan Chen, Changqing Zeng, Fan Liu, Hou-Feng Zheng, Siyang Liu, Hai-Ming Xu, Guo-Bo Chen

## Abstract

Explicitly sharing individual level data in genomics studies has many merits comparing to sharing summary statistics, including more strict QCs, common statistical analyses, relative identification and improved statistical power in GWAS, but it is hampered by privacy or ethical constraints. In this study, we developed ***encG-reg***, a regression approach that can detect relatives of various degrees based on encrypted genomic data, which is immune of ethical constraints. The encryption properties of ***encG-reg*** are based on the random matrix theory by masking the original genotypic matrix without sacrificing precision of individual-level genotype data. We established a connection between the dimension of a random matrix, which masked genotype matrices, and the required precision of a study for encrypted genotype data. ***encG-reg*** has false positive and false negative rates equivalent to sharing original individual level data, and is computationally efficient when searching relatives. We split the UK Biobank into their respective centers, and then encrypted the genotype data. We observed that the relatives estimated using ***encG-reg*** was equivalently accurate with the estimation by KING, which is a widely used software but requires original genotype data. In a more complex application, we launched a finely devised multi-center collaboration across 5 research institutes in China, covering 9 cohorts of 54,092 GWAS samples. ***encG-reg*** again identified true relatives existing across the cohorts with even different ethnic backgrounds and genotypic qualities. Our study clearly demonstrates that encrypted genomic data can be used for data sharing without loss of information or data sharing barrier.

**Author Summary:** Estimating pairwise genetic relatedness within a single cohort is straightforward. However, in practice, related samples are often distributed across different cohorts, making it challenging to estimate inter-cohort relatedness. In this study, we propose a method called encrypted genotype regression (***encG-reg***), which provides an unbiased estimation of inter-cohort relatedness using encrypted genotypes. The genotype matrix of each cohort is masked by a random matrix, which acts similarly to a private key in a cryptographic scheme. This masking process produces encrypted genotypes, which are a projection of the original genotype matrix. We derive the expectation and particularly the sampling variance for ***encG-reg***, the latter involves eighth-order moments calculation. ***encG-reg*** allows us to accurately identify relatedness across cohorts, even for large-scale biobank data. To demonstrate the efficacy of ***encG-reg***, we verified it in a multi-ethnicity UK Biobank dataset comprising 485,158 samples. For this case, we successfully tracked down to the 1st-degree relatedness (such as full sibs and parent-offspring). Furthermore, we used ***encG-reg*** in a collaboration involving 9 Chinese cohorts, encompassing a total of 54,092 samples from 5 genomic centers. It is worth noting that if the number of effective markers is sufficient ***encG-reg*** has the potential to detect even more distant degrees of relatedness beyond what we demonstrated.

## Introduction

Genomic datasets have reached millions of individuals, and are often encapsulated in well-protected cohorts, in which relatives more than often, given increasing genotyped individuals, spread across cohorts and can be identified once the genomic data are compared [1]. Estimating genetic relationship often has clear scientific reasons, such as controlling false positive rates in genome-wide association studies (GWAS) or reducing overfitting in polygenic risk score prediction [2–4]. Social benefits are recently promoted for available individual genomic data such as relativeness testing and forensic genetic genealogy [5]. However, direct-to-consumer (DTC) genetic testing activities along with third-party services pose new privacy and ethical concerns [6]; law enforcement authorities have exploited some of the consumer genomic databases to identify suspects by finding their distant genetic relatives, which has brought privacy concerns to the attention of the general public [7,8]. For regulating forensic genetic genealogy, laws, policies, and privacy-protection techniques are in parallel development [9–11].

The above progress, nevertheless, often requires individual-level data to be shared which may often be beyond the permitted range of data sharing [12]. The encryption methods for genotypes have gone through from the initial one-way cryptographic hashes, to random matrix multiplication, and recently to homomorphic encryption (HE). One-way hashes are leveraged to detect overlapping samples, but it fails if the test genotypes differ, which can be caused by genotyping or imputation errors, even when they are minor [13,14]. Privacy-preserving protocols for multi-center GWAS have been brought to public recently [15–20], on which HE is mostly based. HE provides high precision for results for certain kinds of computational tasks in genetic studies. However, as it is computationally substantial, often one or two orders of magnitude larger than that of the original cost, its application has been limited to small sets of data, at the scale of several hundreds of samples.

In this study, random matrix theory has been adopted to detect relatedness based on our previous study [21]. We developed a novel mitigation strategy called “encrypted genotype regression”, hereby ***encG-reg***, which does not require original genotype data to be shared but is capable of identifying relatedness with highly controllable precision of balanced Type I and Type II error rates. Since only encrypted genotype data is exchanged in performing ***encG-reg***, collaborators from different cohorts are able to minimize their concerns about data confidentiality. We explore the statistical properties of ***encG-reg*** in theory, simulations, and application of 485,158 UK Biobank (UKB) samples of various ethnicity. In a real-world collaboration that includes 5 genomic centers from north to south China (Beijing, Shanghai, Hangzhou, Guangzhou, and Shenzhen) totaling 54,092 genetically diverse samples were genotyped based on different platforms, and intriguing relatedness was identified between cohorts by ***encG-reg***. Privacy-preserving is context-dependent and is still in development under a particular scenario. Throughout this study, when summary information is exchanged, such as allele frequencies and variant positions, we apply the practical guideline in GWAS meta-analysis (GWAMA), which defines a novel strategy we established for data safety.

## Description of the Method

In this section, we will first present an analogous sketch of our thinking. Imagine a Go-like board with dimensions *n*_1_×*n*_2_, where each square contains a particle of size *θ*_*s*_ or *θ*_*l*_, and we assume *θ*_*s*_ < *θ*_*l*_. Additionally, it is often the case that there is a significantly large number of particles of small size *θ*_*s*_ compared to those of larger size *θ*_*l*_. Each particle is imperfect because of the handcraft variance of the particle size. Intuitively, criteria are required if we want to correctly pick a particle of size *θ*_*l*_, which is sampled from a normal distribution 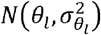, out of many particles of size *θ*_*s*_, which are sampled from 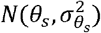. These criteria include the expectation and sampling variance of the particle sizes *θ*_*l*_ we want to pick up, the number of squares on the board (*n*_1_×*n*_2_), the probability that we would accept for incorrectly picking up a particle of size *θ*_*s*_ (Type I error rate α), and the probability that we would miss a real particle of size *θ*_*l*_ (Type II error rate β). All these criteria need to be balanced to find a solution that allows us to distinguish *θ*_*l*_ from *θ*_*s*_ under an acceptable cost, say computational cost and storage cost.

It is evident that the described question fits into the conventional statistical testing scenario, which is “null hypothesis 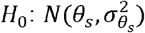 vs alternative hypothesis 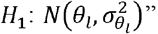. Furthermore, expressions for the minimum size of the testing sample can be given by a power calculator that brings out the required Type I and Type II error rates. In our study, θ refers to the relatedness between a pair of individuals and *m* refers to the number of markers and is related to the sampling variance of relatedness scores. A slightly upscaled concept of *m*, is the effective number of marker *m*_*e*_, which takes into account the squared correlation between *m* markers, and consequently *m*_*e*_ ≤ *m*. Given these considerations, a smaller *m* is preferred to minimize the data cost. Other technical march in **S1 Text** primarily focused on deriving the sampling variance. The variance is approximated using fourth-order moment computation for a fundamental pairwise relatedness estimator, the genetic relationship matrix (GRM). It is further refined through an eighth-order moment computation after multiplying the source genotype matrix by an **S** matrix (*m* rows and *k* columns). Here, the **S** matrix acts analogously to the private key in a cryptographic scheme. The ground truth is that a larger value of *k* facilitates better identification between *θ*_*l*_ and *θ*_*s*_. However, in line with the approach for determining a small yet sufficient value of *m*, we also aim to identify the smallest value of *k* that balances precision and cost. Consequently, the primary objective of this study is to establish a lower bound for *m* (or *m*_*e*_) and *k*. These two values will enable us to determine the minimum conditions required for detecting relatedness.

### Ethic statement

For SBWCH cohort, the protocol and written consent were approved by the Institutional Review Board of BGI (BGI-IRB17088); For CAS cohort, the protocol and written consent were approved by the Institutional Review Board of Beijing Institute of Genomics and Zhongguancun hospital (No.2020H020, No.2021H001, and No.20201229); For ZOC cohort, a written informed consent was obtained from the parents or guardians of the young twins. Ethical approval and DNA data using approval were obtained from the Ethical Committee of Zhongshan Ophthalmic Center; For Fudan cohort, the protocol and written consent were approved by the College of Life Science Fudan University Ethical Review Board; For WBBC cohort, the protocol and written consent were approved by the Westlake University Ethical Review Board.

### Overview of the method

Using SNPs, inter-cohort relatedness for pairs of individuals can be inferred from genetic relationship matrix, which is 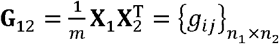. **X**_**1**_ is a matrix of *n*_1_ individuals (rows) and *m* markers (columns), and so is **X**_2._ This GRM definition is identical to equation 9 in Speed and Balding’s review paper for GRM (**X** are standardized by SNP allelic frequencies and its expected sampling variance) [22]. The expectation and variance of *g*_*ij*_. are

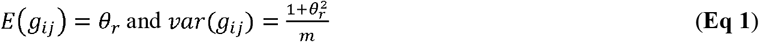

We can express 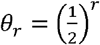 for the *r*^*th*^ degree relatives, so *θ*_*r*_ has the expected values of 1, 0.5, 0.25, and 0.125 for the zero (clonemate), first (full sibs, or parent-offspring), second (half sibs, or grandparent-grandchildren), and third degree (first cousins, or great grandparent-great grandchild) of relatives, respectively. Obviously, when there is an inbred or population structure, or a loop in marriage, the realized value of *θ*_*r*_ covers a continuous range [23].

Let **S** be an *m* × *k* matrix and its entries are independently sampled from *N*(0, σ^2^). 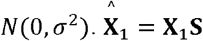 presents an ideal one-way encryption technique in private genetic data sharing, and we call 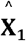 “encrypted genotype”, hereby encG. When 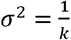, we have 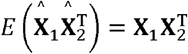, the approximated precision of which relies on the sampling variance of **S** matrix (**S1 Fig**). The relationship is clear: as the value of *k* increases, the approximation approach to the optimum becomes more accurate. In this study, we ask whether there are relatedness existing between **X**_**1**_ and **X**_2_, and how large *k* should be in order to reduce the noise and meanwhile is still able to identify the relatives of certain degree.

Based on encG, it is now trustworthy to construct 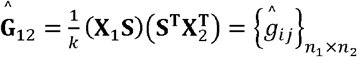, the encrypted GRM. In terms of the matrix element 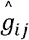, its expectation and variance are 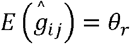 and 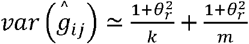, respectively, in which the term 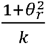 is crept into 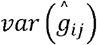 in comparing with its counterpart in **Eq 1**. As SNPs are often in linkage disequilibrium (LD), we introduce the effective number of markers (*m*_*e*_), which is a population parameter engaged in various genetic analyses [24]. The variances of *g*_*ij*_ and 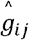 then become 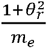 and 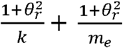, respectively.

### encG regression (encG-reg)

Another interpretation of encGRM is from the perspective of linear regression, which we regress the *r*^*th*^ row of 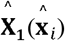 against the *j*^*th*^ row of 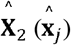 to estimate the relatedness. We call this procedure the encG regression (***encG-reg***). The slope *g*_*ij*_ of a simple regression model 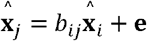 indicates the relatedness score between these two0020individuals. The expectation and the sampling variance of 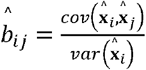 can be approximated by

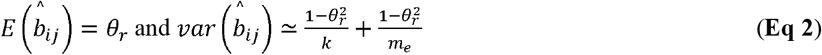

Compared with encGRM, encG-reg generates a smaller sampling variance and improves statistical power in identifying relatives.

### Minimum numbers of *m*_*e*_ and *k*

Given a pair of individuals **I**) whose relatedness is estimated by GRM and follows the distribution of 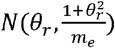, we ask how to identify them from unrelated pairs with a distribution of 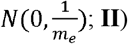 whose relatedness is estimated by ***encG-reg*** and follows the distribution of 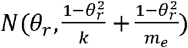, we ask how to differentiate them from unrelated individual pairs as sampled from 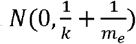. This question is analogous to the conventional pattern recognition, which can be solved under the power calculation in the statistical test framework for null verse alternative hypotheses. We consequently need to determine two key parameters. **I**) the effective number of markers, *m*_*e*_, a population statistic that sets the resolution of GRM itself in detecting relatives. **II**) the column number of the random matrix, *k*, an iteration dimension that sets the precision of ***encG-reg***. To determine *m*_*e*_ and *k*, upon Type I error rate (α, false positive rate as aforementioned) and Type II error rate (β, false negative rate), *m*_*e*_ should satisfy the below inequality

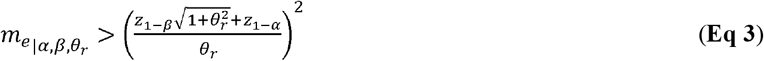

Similar to *m*_*e*_, the minimum number of *k* is also determined by a certain Type I and Type II error rates, the degree of relatives to be detected, as well as the parameter *m*_*e*_. *k* should follow the below inequality

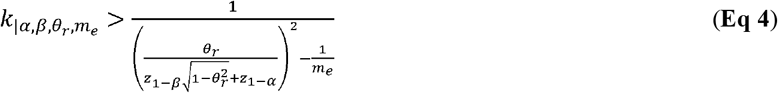

In particular, α should be under experiment-wise control, say after Bonferroni correction, and consequently upon the total comparisons 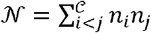, where there are 𝒞 cohorts and *n*_**1**_ samples in cohort *i*. Of note, **Eq 3** gives a lower bound of the number of markers, while in practice we often have genome-wide SNPs in surplus, such as the case in the UKB example below. As a larger *m*_*e*_ leads to a smaller *k*, it is upon the data to choose a large *m*_*e*_ but a small *k*, or a minimum *m*_*e*_ but a large *k*. **Fig 1** provides a phenomenological illustration of how *m*_*e*_ and *k* are weaved together. A simulation R code for **Fig 1** can be found at https://github.com/qixininin/encG-reg/blob/main/1-Simulations/Figure1-resolution.R.

**Fig 1.**
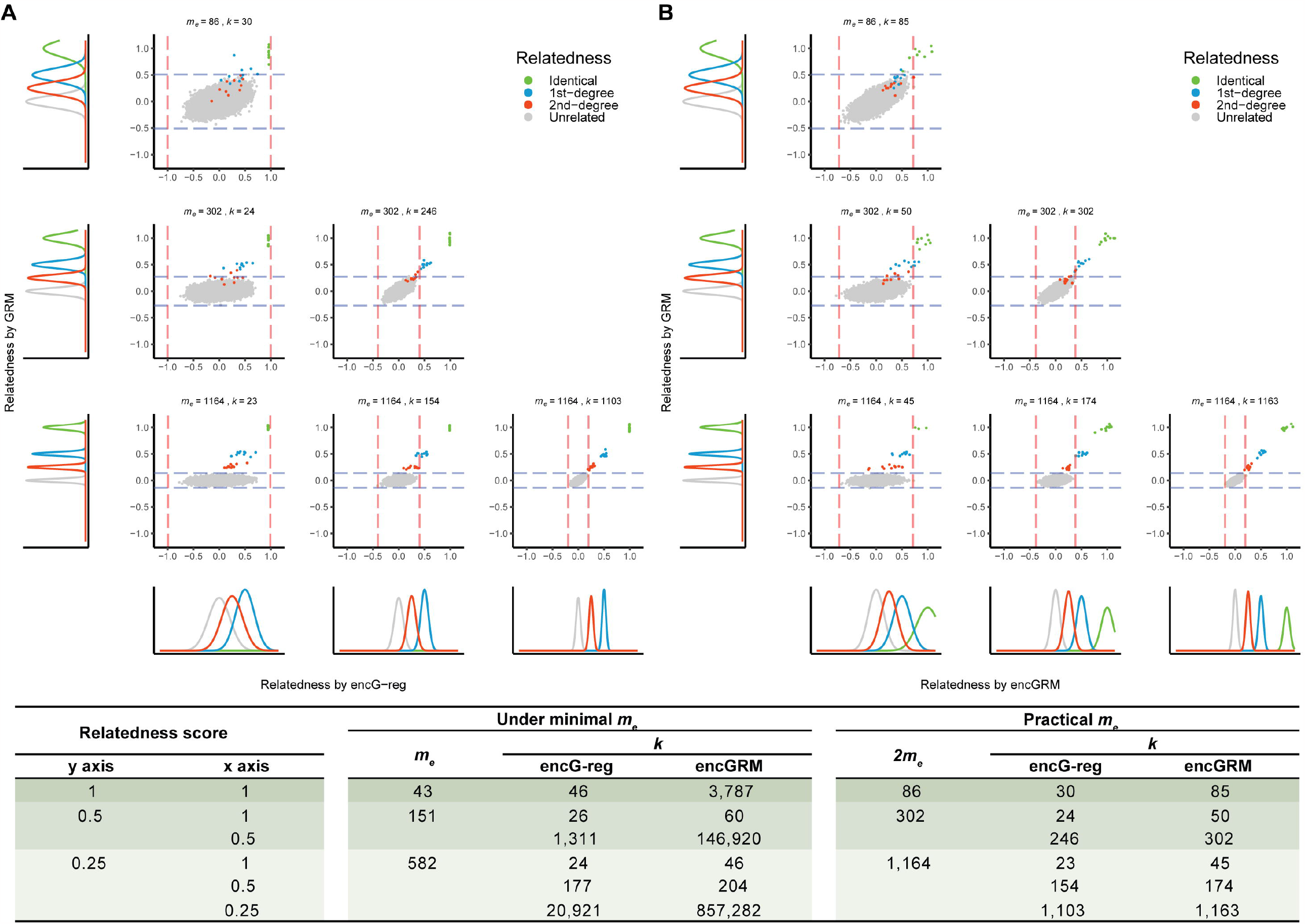
Resolution for varying relatedness using GRM, encGRM and encG-reg. The figure shows the resolution for detecting relatives or overlapping samples with respect to varying number of markers at every row (for better illustration *m*_*e*_ was twice that of **Eq 3**) and the degree of relatives to be detected (*r*=0, 1, and 2). The y axis is the relatedness calculated from GRM and the x axis is the estimated relatedness calculated from ***encG-reg*** (**A**) and encGRM (**B**). Each point represents an individual pair between cohort 1 and cohort 2 (there are 200 × 200=40,000 pairs in total), given the simulated relatedness. The dotted line indicates the 95% confidence interval of the relatedness directly estimated from the original genotype (blue) and the encrypted genotype (red). The table provides how *m* and *k* are estimated. The columns “under minimal *m*” provide benchmark for a parameter, and it is practically to choose 2 × *m*_*e*_ and then estimate *k* as shown under the column “practical *m*_*e*_”.

The conceptual layout of the method is as described above. The technical details of sampling variance at **Eq 1-2** and statistical power calculation for **Eq 3-4** can be found in **S1 Text** and the corresponding annotations are given in **S1 Table**. The properties of all three methods, including GRM, encGRM, and ***encG-reg***, are summarized in **Table 1**. For a pair of cohorts of sample *n*_1_ and *n*_2_, the computational time complexity of ***encG-reg*** is about 𝒪((*n*_1_ + *n*_2_)*mk + n*_1_ *n*_2_*k*): the first term occurs at the local site of each cohort and the second term occurs at an entrusted computational server for a pair of cohorts. Furthermore, after local encryption by multiplication of **S** the size of the 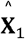 that is sent to the central analyst is of dimension *n*_1_ × *k*, which approximately represents the space complexity. Although the value of *m* and *k* are as determined by **Eq 3** and **Eq 4**, but in practice we may pick a slightly larger *m*, say 2 *m*, so as to balance time complexity and space complexity.

**Table 1.** Statistical properties of conventional genetic relationship matrix (GRM), encrypted GRM (encGRM), and encG regression (encG-reg).

|  | GRM | encGRM | encG-reg |
| --- | --- | --- | --- |
| **Matrix form** | $\mathbf{G}_{12} = \frac{\mathbf{X}_1 \mathbf{X}_2^T}{m} = \{g_{ij}\}_{n_1 \times n_2}$ | $\hat{\mathbf{G}}_{12} = \frac{(\mathbf{X}_1 \mathbf{S})(\mathbf{S}^T \mathbf{X}_2^T)}{k} = \{\hat{g}_{ij}\}_{n_1 \times n_2}$ | $\hat{\mathbf{B}}_{12} = \{\hat{b}_{ij}\}_{n_1 \times n_2}$ |
| **Expectation** | $E(g_{ij}) = \theta_r$ | $E(\hat{g}_{ij}) = \theta_r$ | $E(\hat{b}_{ij}) = \theta_r$ |
| **Variance** | $var(g_{ij}) = \frac{1 + \theta_r^2}{m_e}$ | $var(\hat{g}_{ij}) \approx \frac{1 + \theta_r^2}{k} + \frac{1 + \theta_r^2}{m_e}$ | $var(\hat{b}_{ij}) \approx \frac{1 - \theta_r^2}{k} + \frac{1 - \theta_r^2}{m_e}$ |
| **Time complexity** | $\mathcal{O}(n_1 n_2 m)$ | $\mathcal{O}((n_1 + n_2)mk + n_1 n_2 k)$ | $\mathcal{O}((n_1 + n_2)mk + n_1 n_2 k)$ |
| **Space complexity<sup>a</sup>** | | $\mathcal{O}(n_i k)$ | $\mathcal{O}(n_i k)$ |

**Table notes:** This table compares the definition and properties, such as expectation, variance and time complexity between three important matrices, GRM, encGRM, and *encG-reg*, mentioned in the article. Note that, each element of the random matrix $\mathbf{S}_{m \times k} = \{s_{ij}\}$ follows a normal distribution $N(0, 1/m)$ ; $m$ is the number of markers, also represents the number of rows for random matrix $\mathbf{S}$ ; $k$ is the number of columns for random matrix $\mathbf{S}$ ; $m_e$ is the number of effective markers, an advanced concept that takes the squared correlation between markers into account; $r$ is the degree of a pair of relatives and $\theta_r$ is their relatedness score; $n_1$ and $n_2$ are the sample sizes of two cohorts.
<sup>a</sup>Space complexity refers to the dimension of $\hat{\mathbf{X}}_i = \mathbf{X}_i \mathbf{S}$ , which should be transported to the central analyst.

After the assembly of cohorts, there are options for choosing SNPs upon the experimental design. An exhaustive design denotes the use of intersected SNPs between each pair of cohorts, thus a specific random matrix will be shared with each pair of cohorts. Given 𝒞 cohorts, there are 𝒞(𝒞 − 1)/2**S** matrices generated and each cohort is likely to receive 𝒞 − 1 different **S** matrices that matches to 𝒞 − 1 cohorts. Adopting exhaustive design is possible to maximize the statistical power with the maximized number of SNPs, but the computational, as well as communicational, efforts may overwhelm the organization of a study. In contrast, a parsimony design denotes the use of intersected SNPs among all assembled cohorts, as long as the number of SNPs satisfies the resolution in **Eq 3** and **Eq 4**. Exhaustive design and parsimony design are both validated in the 19 UKB cohorts, each of which had sample sizes greater than 10,000, and parsimony design is further tested in the real world for 9 Chinese cohorts in this study.

### Protocol for encG-reg for biobank-scale application

We now sketch a detailed technical protocol with security concern for ***encG-reg*** into four steps and two interactions. For the four steps, steps 1 and 3 are performed by each collaborator, and steps 2 and 4 are performed by a central analyst (**Fig 2A**). For the two interactions, exchanged information, possible attacks, and corresponding preventative strategies are given in examples (**Fig 2B**). We have provided comprehensive details and practical commands for each step at https://github.com/qixininin/encG-reg. The repository includes two main folders: “Simulation” and “Protocol”. The “Simulation” folder includes code and resources for all simulations in this study. The “Protocol” folder contains a user-friendly protocol that outlines the step-by-step process for a group of collaborators using ***encG-reg***. These commands and scripts have been utilized during interactions with our collaborators in multi-center Chinese datasets application.

**Fig 2.**
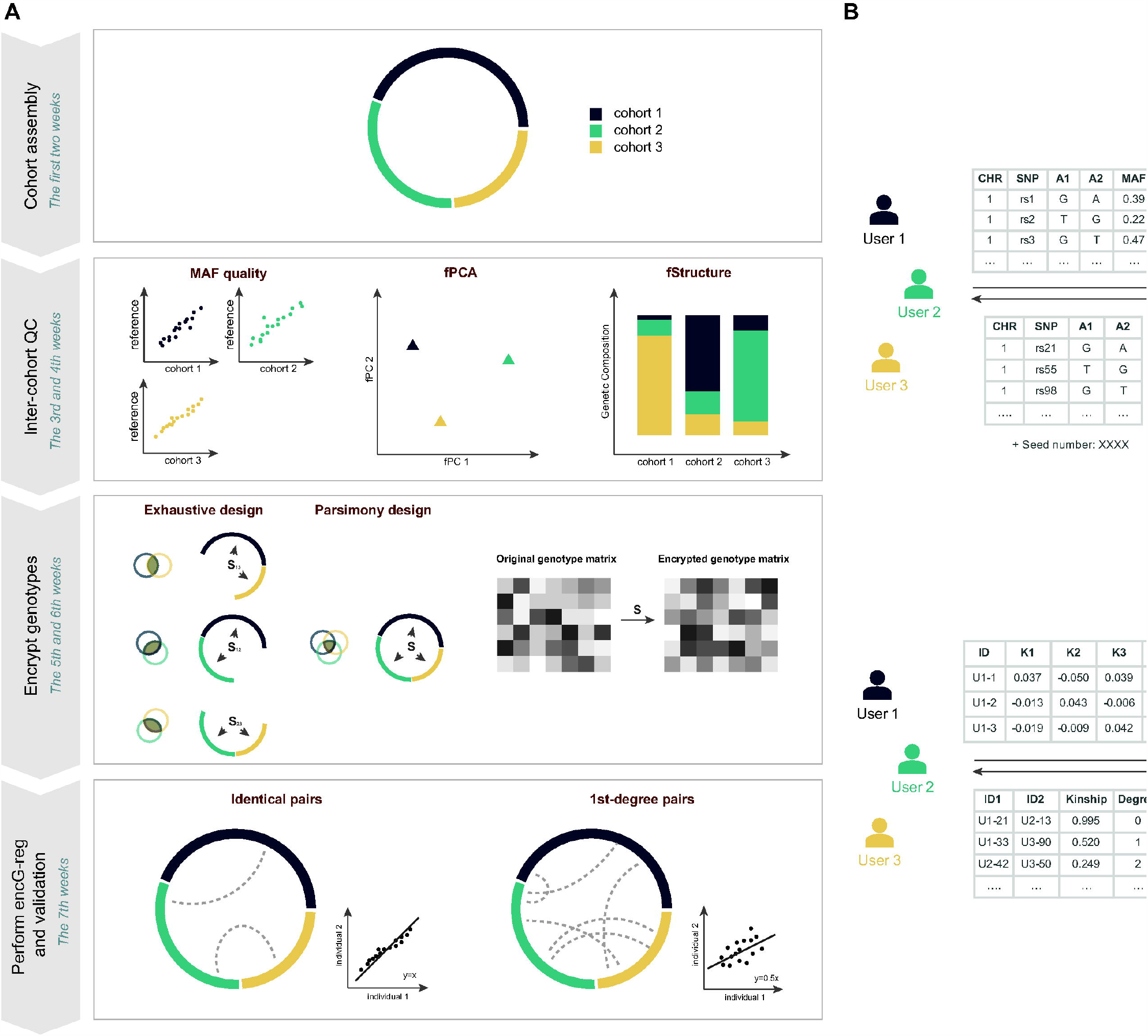
Workflow of encG-reg and its practical timeline as exercised in Chinese cohorts. The mathematical details of ***encG-reg*** are simply algebraic, but its inter-cohort implementation involves coordination. (**A**) We illustrate its key steps, the time cost of which was adapted from the present exercise for 9 Chinese datasets (here simplified as three cohorts). **Cohort assembly**: It took us about a week to call and got positive responses from our collaborators (See **Table 3**), who agreed with our research plan. **Inter-cohort QC**: we received allele frequencies reports from each cohort and started to implement inter-cohort QC according to “geo-geno” analysis (see **Fig 6**). This step took about two weeks. **Encrypt genotypes**: upon the choice of the exercise, it could be exhaustive design (see UKB example), which may maximize the statistical power but with increased logistics such as generating pairwise **S**_*ij*_; in the Chinese cohorts study we used parsimony design, and generated a unique **S** given 500 SNPs that were chosen from the 7,009 common SNPs. It took about a week to determine the number of SNPs and the dimension of *k* according to **Eq 3** and **4**, and to evaluate the effective number of markers. **Perform encG-reg and validation**: we conducted inter-cohort ***encG-reg*** and validated the results (see **Fig 7** and **Table 4**). It took one week. (**B**) Two interactions between data owners and central analyst, including example data for exchange and possible attacks and corresponding preventative strategies.

#### Step 1 Cohort assembly and intra-cohort quality controls

Basic intra-cohort QCs should be conducted. Summary information such as SNP ID, reference allele, and its frequency are then requested by the central analyst.

#### Step 2 Inter-cohort quality controls and parameter setup

Using the “geo-geno” relationship often observed in genetic data [25,26], we suggest two inter-cohort QCs. One is called frequency-principal component analysis (fPCA) and another is called fStructure. The technical details of the employed fPCA and fStructure methods can be found in our previous study [21] and is described in the following subsection. The central analyst determines *m* and *k* by **Eq 3** and **Eq 4** based on survived SNPs and passes parameter information to each collaborator along with a SNP list.

#### Step 3 Encrypt genotype matrix

The *m*-by-*k* random matrix, or matrices when an exhaustive design is chosen, is generated and sent to each collaborator. As a positive control, reference samples will be merged into each cohort. Genotype encryption is realized by the matrix multiplication between the standardized genotype matrix and **S**.

#### Step 4 Perform encG-reg

Inter-cohort computing for relatedness will be conducted by the central analyst. A successful implementation will lead to at least positive controls consistently identified as inter-cohort “overlap” and if possible, various sporadic relatedness.

In the above steps, there are two interactions between collaborators and the central analyst:

#### The first interaction

Collaborators send over a list of variants including their allele frequencies. After doing variants selection, central analyst returns a list of intersected variants, together with a randomly generated seed. Re-identifications based on allele-frequency may occur, but a suggested choice of common variants (MAF>0.05) can mostly dispel these misgivings.

#### The second interaction

Collaborators send over a matrix of encrypted genotypes. After performing ***encG-reg*** between each two pairs of cohorts, the central analyst returns identified relatedness or returns relatedness scores directly, based on pre-agreed requests. PCA-attack based on encrypted genotypes may occur, if the correlation structure of variants being approximated in a proper reference population, but one straightforward defense is to use variants that are in linkage equilibrium, and it ensures that the correlation matrix closely approximates to the identity matrix.

### Cohort-level quality control using fPCA and fStructure

In this study, we use fPCA and fStructure to examine the data quality at the cohort-level using summary statistics. Both fPCA and fStructure has been previously used in The Genetic Investigation of Anthropometric Traits (GIANT) Consortium [21], and GWAMA for educational attainment [27]. fPCA is a principal component analysis based on summary data at the population level, rather than individual-level data. It uses the allele frequency of common markers across all cohorts, which is constructed into matrix **P** = {*p*_*ij*_} _𝒞 × ℳ_ Here, 𝒞 represents the number of cohorts and ℳ represents the number of common markers, while *p*_*ij*_ denotes the allele frequency for the *j*^*th*^ marker in the *i*^*th*^ cohort. Using the differences in marker frequencies, fPCA can effectively capture the population structure in PC1 and PC2. fStructure explores the genetic composition of the target cohorts by comparing with reference populations using *F*_*st*_. Again, *F*_*st*_ is calculated based on allele frequency of common markers among all cohorts and the reference populations. Suppose there are 𝒞 cohorts and three reference population. The average *F*_*st*_ using common markers (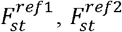, and 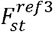) between *i*^th^ cohort and the given reference populations are calculated. Finally, a bar plot is employed to show the ratio of 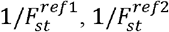, and 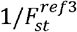, providing a clear visualization of the genetic composition of the testing cohorts against the reference populations.

### Calculate the number of effective markers

The corresponding *m*_*e*_ will be estimated from, 1KG-EUR and 1KG-CHN as the reference populations for validation in the UKB cohorts and the Chinese cohorts, respectively. According to its definition, 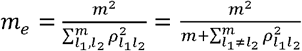, in which *ρ*^2^ is the squared Pearson’s correlation for a pair of SNPs, and *m*_*e*_ can be empirically estimated as 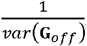, where **G**_*off*_ denotes the off-diagonal elements of GRM [24,28,29]. 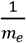 describes the global LD for all the included SNPs. For more technical details about *m*_*e*_, please refer to Huang et al [30]. Since *m*_*e*_ is asymptotically distributed as 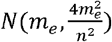 according to the Delta method, the sampling variance of *m*_*e*_ is negligible as long as the studying populations are from similar ancestries, such as the case for Manchester and Oxford cohorts in UKB and the Chinese datasets employed in this study (**S2 Table**). For a single-ethnicity population, when SNPs are randomly sampled from the genome, with *m* < 50,000, *m*_*e*_ is approximately equal to *m*, as demonstrated in Chinese cohorts. In the case of a multi-ethnicity population like UKB, the use of ***encG-reg*** is when SNPs of ethnicity-insensitive frequencies are employed.

## Verification and Comparison

### Simulation validation

For a conceptional exploration, we illustrated how *m* and *k* would affect the identification of various relatedness. In this case, we ignored the difference between *m* and *m*_e_ because SNPs were generated independently. We simulated 200 individuals each for cohort 1 and cohort 2 (*n*_1_ = *n*_2_ = 200). Between cohort 1 and cohort 2, we generated 10 pairs of related samples at a variety of relatedness, i.e., zero-degree/identical, 1st-degree, and 2nd-degree relatives, respectively. For better illustration, we set the desired number of markers (*m*) twice as given by **Eq 3** and the corresponding size of *k* as given by **Eq 4** at the experiment-wise Type I error rate of 0.05 (α =0.05/40,000) and Type II error rate of 0.1 – equivalent to 90% statistical power. We simulated individual-level genotype matrices with the dimension of *n*_1_ × *m* and *n*_2_ × *m* and the encrypted genotype matrices with the dimension of *n*_1_ × *k* and *n*_2_ × *k*. Relatedness scores for GRM, encGRM, and ***encG-reg*** were calculated accordingly and theoretical distributions were derived under the assumption of multivariate distribution for each degree of relatedness. **Fig 1** showed that for ***encG-reg***, in each scenario, sufficient *k* was able to detect a certain degree of relatedness as long as *m* could support. Compared with encGRM, ***encG-reg*** had a smaller variance and consequently a larger statistical power in detecting relatives.

We then evaluated the properties and performance of ***encG-reg***, GRM, and encGRM in more details. We first validated the derived variances of GRM, encGRM, and ***encG-reg*** (as summarized in **Table 1**). 1,000 pairs of relatives were separated in cohort 1 and cohort 2. *m* = 1,000, 1,250, 1,500, 1,750, and 2,000 independent markers were simulated, and their MAF was sampled from a uniform distribution *U*(0.05,0.5). Genotype matrices from two cohorts were encrypted by the same *m* × *k* random matrix **S**, whose elements drew from a normal distribution 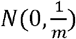). We set *k* to be 1,000, 2,000, 3,000, 4,000, and 5,000, respectively. Both the original and the encrypted genotype matrices were standardized based on the description for the three methods. Observed and theoretical variances were examined among four different degrees of relatedness (identical, 1st-degree, 2nd-degree, and 3rd-degree). The estimated sampling variances of GRM, encGRM and ***encG-reg*** matched with the theoretical variance at each level of relatedness (**Fig 3**).

**Fig 3.**
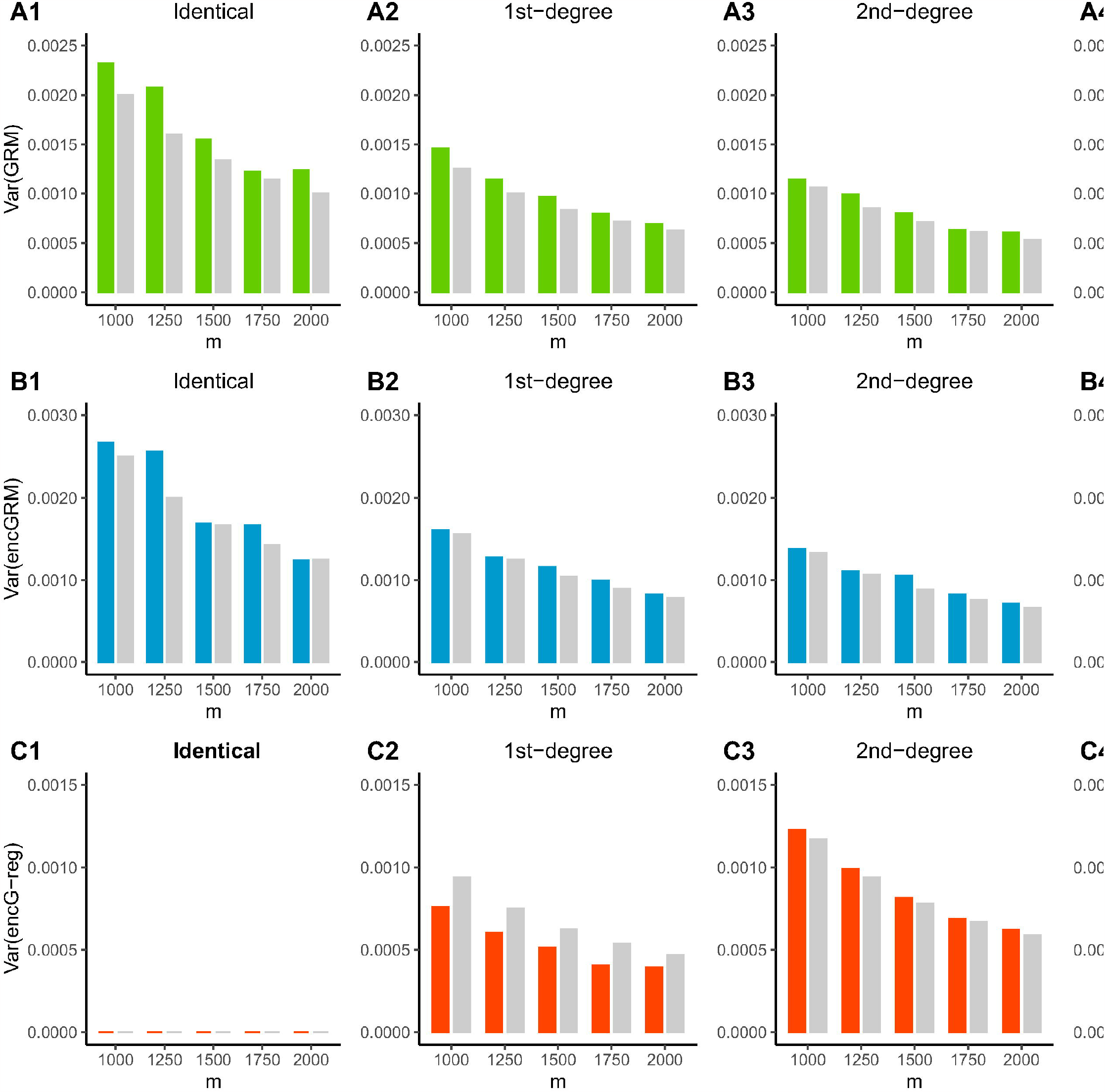
Sampling variance of GRM, encGRM and encG-reg in simulations. The observed and theoretical sampling variance of GRM (**A1-A4**), encGRM (**B1-B4**) and ***encG-reg*** (**C1-C4**) are given in bar plots. Individual genotypes are simulated with *m* = 1,000, 1,250, 1,500, 1,750, and 2,000 independent markers. A total number of *n*_1 =_ *n*_2_ = 1,000 pairs of relatives are simulated under each different levels of relatedness (*r* = 0, 1, 2, and 3). As for the encryption, the column number of random matrices are *k* = 4,000, 5,000, 6,000, 7,000, and 8,000 correspondingly.

### Multi-ethnic samples validation: UKB in exhaustive and parsimony design

Both exhaustive and parsimony designs were conducted to validate ***encG-reg*** on 485,158 UKB multi-ethnicity samples from 19 assessment centers with a sample size greater than 10,000 (**S3 Table**), resulting in a total of 110,713,926,381 inter-cohort individual pairs. As the 485,158 UKB samples consist of 94.23% Whites, 1.94% Asian or Asian British, 1.57% Black or Black British, and 2.27% “Other” or “Unknown”, it is an ideal dataset to validate whether ***encG-reg*** is good to handle diversified samples. Identical/twins, 1st-degree and 2nd-degree relatedness were aimed to be detected by KING-robust (“the rule of thumb”) using the real genotypes and by ***encG-reg*** using the encrypted genotypes, respectively. We conducted QC on the 784,256 chip SNPs within the 19 cohorts, and the inclusion criteria for autosome SNPs were: (1) minor allele frequency (MAF) > 0.01; (2) Hardy-Weinberg equilibrium (HWE) test *p*-value > 1e-7; and (3) locus missingness < 0.05. An averaging number of 578,543 SNPs survived from 19 cohorts. In addition, taking account of the multi-ethnicity nature of UKB samples, only SNPs of ethnicity-insensitive frequency, which have indifferent allele frequencies statistically, were included. Therefore, we selected ethnicity-insensitive SNPs based on three reference population representing European (99 CEU and 107 TSI samples), African (107 YRI samples), or East Asian (103 CHB and 105 CHS samples) background in 1000 Genome Project. We first performed SNP quality control on three reference population, using the following inclusion criteria: (1) minor allele frequency (MAF) > 0.01; (2) Hardy-Weinberg equilibrium (HWE) test *p*-value > 1e-7; and (3) locus missingness < 0.05. Next, we conducted association studies on two reference populations at a time, selecting SNPs with a *p*-value greater than 0.05 (considered insignificant). Once insignificant SNPs were identified between each pair of reference population, we took the intersection between those SNPs to establish the ethnicity-insensitive SNP pool, which contained a total of 299,835 SNPs. These SNPs exhibit consistent allele frequency across different ethnic backgrounds. For more details see **S1 Text**.

We adopted both exhaustive design and parsimony design in this UKB validation. In the exhaustive design, intersected SNPs were selected between each pair of cohorts, the average number of intersected SNPs was 556,929 and the average number of ethnicity-insensitive SNPs after taking intersection with the ethnicity-insensitive pool was 13,157. In the parsimony design, a total number of 12,858 intersected and also ethnicity-insensitive SNPs among all 19 cohorts were selected. The numbers of ethnicity-insensitive SNPs intersected between each pair of UKB cohorts in exhaustive design were all given in **S4 Table**. The number of *k* for ***encG-reg*** was estimated by **Eq 4** at a Type I error rate of 0.05 and a Type II error rate of 0.1. To note that, experiment-wise Bonferroni correction is based on the number of paired samples between every two cohorts (𝒩_*ij*_ = *n*_*i*_ *n*_*j*_) for exhaustive design and the total number of paired samples among all cohorts 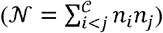 for parsimony design.

### Performance of encG-reg in two UKB cohorts

We investigated more details of ***encG-reg*** at two assessment centers in Manchester (11,502 individuals) and Oxford (12,260 individuals) from UKB white British, which included over 140 million comparisons. We randomly sampled SNPs with different ranges of MAF (0.01 to 0.05, 0.05 to 0.15, 0.15 to 0.25, 0.25 to 0.35, 0.35 to 0.5, and a broad range of 0.05 to 0.5) so as to compare the performance of ***encG-reg*** and KING. According to the minimum number of *m*_*e*_ and *k* at the experiment-wise Type I error rate of 0.05 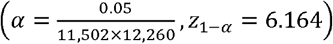 and Type II error rate of 0.1 (z_1 − β_) based on **Eq 3** and **Eq 4** (**Table 2**), the minimum requirement for *m*_*e*_ was 283 and 1,104 for detecting 1st-degree and 2nd-degree relatedness, respectively. However, since it is *m* rather than *m*_*e*_ that can be directly determined and interacted with the data, we suggested and empirically chose *m* as twice the minimum number of *m*_*e*_, in order to ensure that the practical derived *m*_*e*_ satisfies **Eq 3**. We randomly selected 566 SNPs (*m*_*e*_ = 566, θ_1_ = 0.45) and 2,209 SNPs (*m*_*e*_ = 2,023, θ_2_ = 0.225) for detecting 1st-degree and 2nd-degree relatedness, and the corresponding *k* were 494 and 2,342, respectively. Against possible noise that may rust statistical power, we also increased *k* to 1.2*k* and denoted it as encG-reg+. The average relatedness score, standard deviation, and statistical power were calculated for each detected relative pair after resampling SNPs 100 times.

**Table 2.** The minimum number of *m*_*e*_ and *k* for identifying different relatedness between two UKB cohorts Manchester and Oxford.

| Relatedness | Minimum $m_e$ (Eq 3) | Suggested | | |
| --- | --- | --- | --- | --- |
| | | $m$ (empirical $m_e$) | $k$ (Eq 4) | $1.2 \times k$ |
| Identical ($\theta = 1$) | 77 | 154 (157) | 87 | 105 |
| 1st-degree ($\theta = 0.5$) | 283 | 566 (566) | 494 | 593 |
| 2nd-degree ($\theta = 0.25$) | 1,105 | 2,209 (2,023) | 2,342 | 2,811 |

**Table notes:** We used Manchester (11,502 individuals) and Oxford (12,260 individuals) from UKB white British, totaling $11,502 \times 12,260 = 141,014,520$ pairs. The minimum number of $m_e$ for detecting identical, 1st-degree, and 2nd-degree relatedness are calculated from **Eq 3** at the experiment-wise Type I error rate of 0.05 ( $\alpha = 0.05/141,014,520$ ) and Type II error rate of 0.1. In practice, the suggested number of markers ( $m$ ) here is twice that of the minimum number of $m_e$ because it is a surplus of SNPs. The empirical $m_e$ is estimated from 1KG-EUR. The minimum number of $k$ is calculated from **Eq 4** given empirical $m_e$ . The column of “ $1.2 \times k$ ” is suggested for improved statistical power.

Out of the 11,502 x12,260 = 141,014,520 pairs of inter-cohort individuals, 17 pairs of so-called 1st-degree and 2 pairs of 2nd-degree relatives were found using overall QCed SNPs by KING. The bar plots in **Fig 4** compared relatedness scores of the known 1st-degree (*m*_*e*_ = 566, *k* = 494) and 2nd-degree (*m*_*e*_ = 2,023, k = 2,342) relatives, estimated by KING, GRM, ***encG-reg***, and ***encG-reg+*** (using 1.2*k*). In general, ***encG-reg*** and ***encG-reg+***, still showed very similar estimations of relatedness score compared with KING. When SNPs were sampled with MAFs between 0.05 and 0.5, the average statistical power reached 0.9 and 0.95 for detecting 1st-degree relatedness by ***encG-reg*** and ***encG-reg+***. The overall statistical power was proportional to the MAF; when the MAF of the sampled SNPs was less than 0.05, the statistical power of ***encG-reg*** was still close to our theoretical benchmark. In a more refined scope, using the conditional binomial distribution, our analytical result showed that the sampling variance of GRM was proportional to 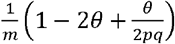 (**S1 Text**). It was noticeable that larger MAFs could lead to a smaller variance of GRM score (**S2 Fig)**, which further resulted in a smaller variance and a higher power of detecting relatives for encGRM and ***encG-reg***. This result is consistent with how MAF affects the statistical power in UKB Manchester and Oxford cohorts.

**Fig 4.**
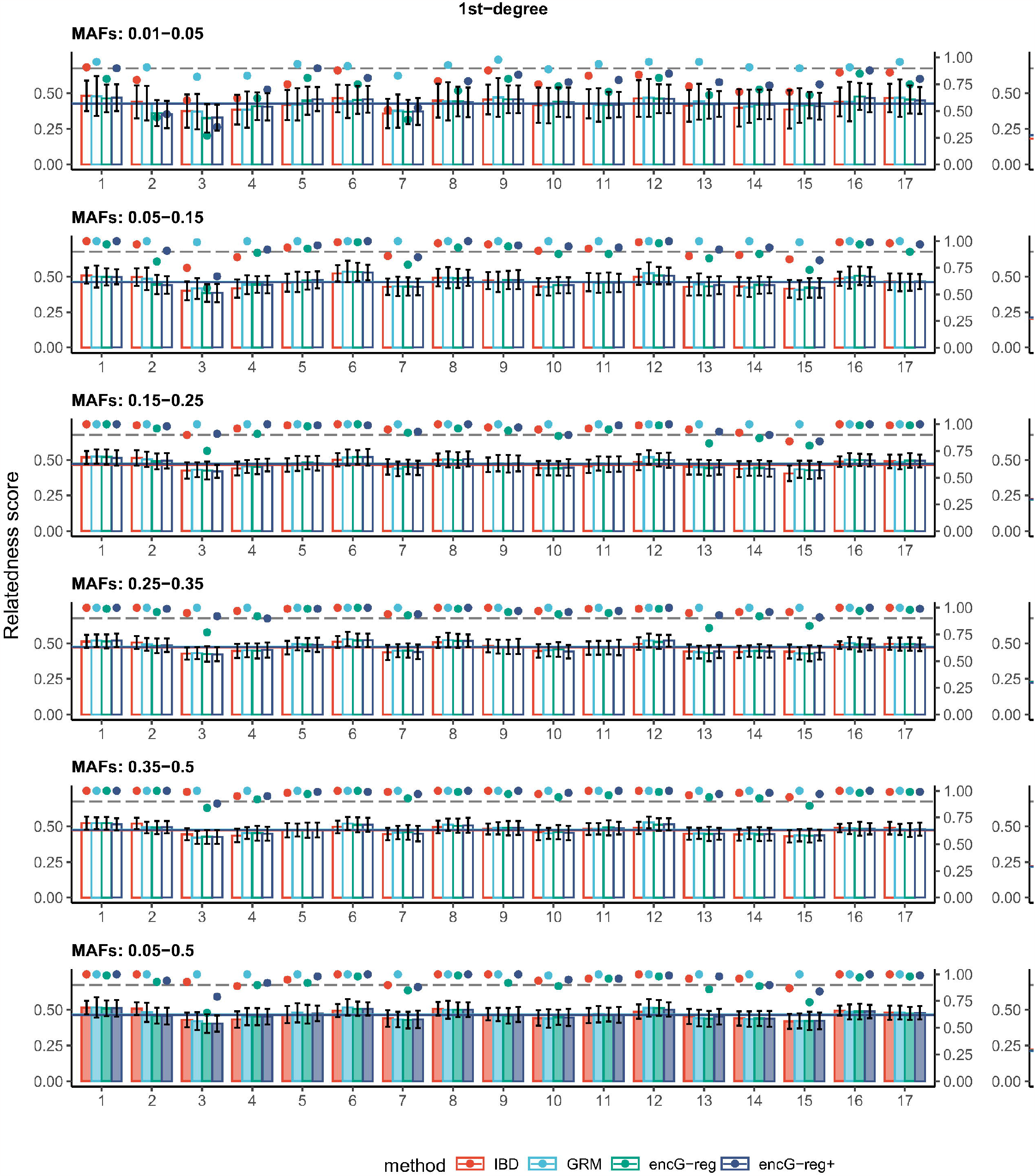
Influence of minor allele frequencies in detecting relatives in Manchester and Oxford cohorts. The bar plots provide a comparison of relatedness scores for the known 1st-degree and 2nd-degree relatives estimated by KING, GRM, ***encG-reg***, and ***encG-reg+*** at two representative assessment centers (Manchester and Oxford). For each assessment centers, 566 and 2,209 SNPs were randomly selected within specific MAF ranges: 0.01 to 0.05, 0.05 to 0.15, 0.15 to 0.25, 0.25 to 0.35, 0.35 to 0.5, and 0.05 to 0.5. Here, ***encG-reg***+ denotes the use of 1.2-fold of the minimum number of *k* and IBD denotes twice the relatedness score estimated by KING. After resampling SNPs 100 times, the average GRM score, standard deviation, and statistical power were calculated for each detected relative-pair. The grey dashed line indicates the expected statistical power of 0.9. The solid colored lines indicate the average relatedness scores for certain degrees as estimated by the four methods. 17 pairs of so-called 1st-degree and 2 pairs of 2nd-degree relatives were approved using overall SNPs by KING.

### Performance of encG-reg in UKB

We verified the exhaustive design of ***encG-reg*** in 19 UKB cohorts (totaling over 100 billion inter-cohort individual pairs) by comparing with the results from KING up to the 2nd-degree relatedness (**Fig 5A**). The average number of intersected SNPs between every two pairs of cohorts was 13,157. The same 38 pairs of identical samples (monozygotic twins in this case) were detected by KING and ***encG-reg***; 7,965, and 6,632 pairs of 1st-degree and 2nd-degree relatedness were inferred by KING, comparing to 7,913 and 7,022 by ***encG-reg***, respectively. It could be seen that ***encG-reg*** was quite comparable to KING in practice. Based on individual ID and their recorded ethnicity, consistent relatedness scores were estimated by KING and ***encG-reg*** (**Fig 5B-5D**). Combining geographic distance between 19 cohorts, we discovered that more relatives were detected between adjacent assessment centers, such as Manchester and Bury, Newcastle and Middlesbrough, and Leeds and Sheffield. Besides, consistent numbers of relatedness were inferred by the parsimony design of ***encG-reg*** (**S5 Table**). The decrease in the number of the detected 2nd-degree relatedness for parsimony design was possibly due to the smaller experiment-wise Type I error rate and thus a more stringent threshold.

**Fig 5.**
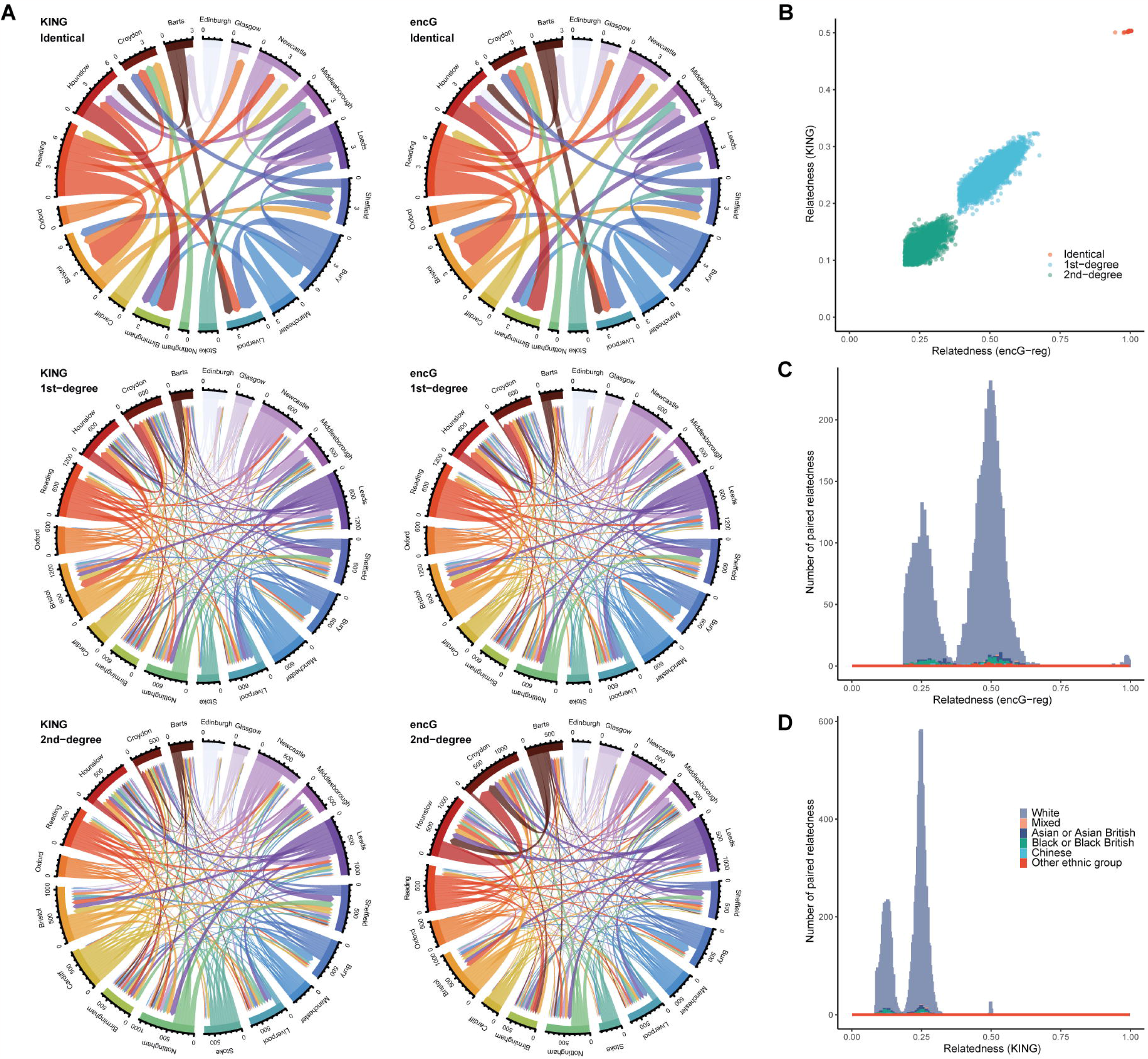
Resolution for detecting relatives in UKB cohorts by KING and encG-reg under exhaustive design. (**A**) Chord diagrams show the number of inter-cohort identical/twins, 1st-degree, and 2nd-degree relatedness across 19 UKB assessments with over 10,000 samples. Relatedness was detected and compared between KING and ***encG-reg*** under an exhaustive design, encompassing a total of 171 inter-cohort analyses. In each chord plot, the length of the side edge is proportional to the count of detected relatives between the focal cohort and other cohorts. (**B**) The scatter plot shows the estimated relatedness score by KING and ***encG-reg*** for inter-cohort relative pairs, including identical, 1st-degree, and 2nd-degree pairs. (**C**) The histogram shows the distribution of relatedness scores estimated by ***encG-reg***. (**D**) The histogram shows the distribution of relatedness scores estimated by KING.

As aforementioned the computational time complexity was 𝒪 (*n*_l_ + *n*_2_)*mk* + *n*_l_*n*_2_*k*). For the example of 19 UKB cohorts, with an average cohort size of *n* =25,537, an average of *m* =13,157 intersected ethnicity-insensitive markers, and an average of *k* =1,381 columns for random matrix, the average time required for each pair-wise ***encG-reg*** computation was 9.682 ± 2.700 minutes (**S4 Table**). The computations were performed using one thread on an Intel(R) Xeon(R) CPU E7-4870 @ 2.40GHz. The average storage space of data, which was supposed to be transported to the center analyst but not for the UKB in-house demonstration, was proportional to *n* × *k*. Both the computational time and the storage space was affordable even for biobank-scale data such as the UKB.

## Applications

### 9 multi-center Chinese datasets

We launched a national-scale application for ***encG-reg*** in 9 Chinese datasets under the parsimony design to avoid possible computational and communicational costs. Four out of nine datasets were publicly available, while the remaining datasets were recruited from 5 research centers, located in from north to south China, including Beijing, Shanghai, Hangzhou, Guangzhou, and Shenzhen (**Table 3**). Serving as a proof-of-concept and brief validation of ***encG-reg*** in civilian and complex environments, collaboration was organized to detect identical or 1st-degree relatedness samples but without revealing personal medical information.

**Table 3.**
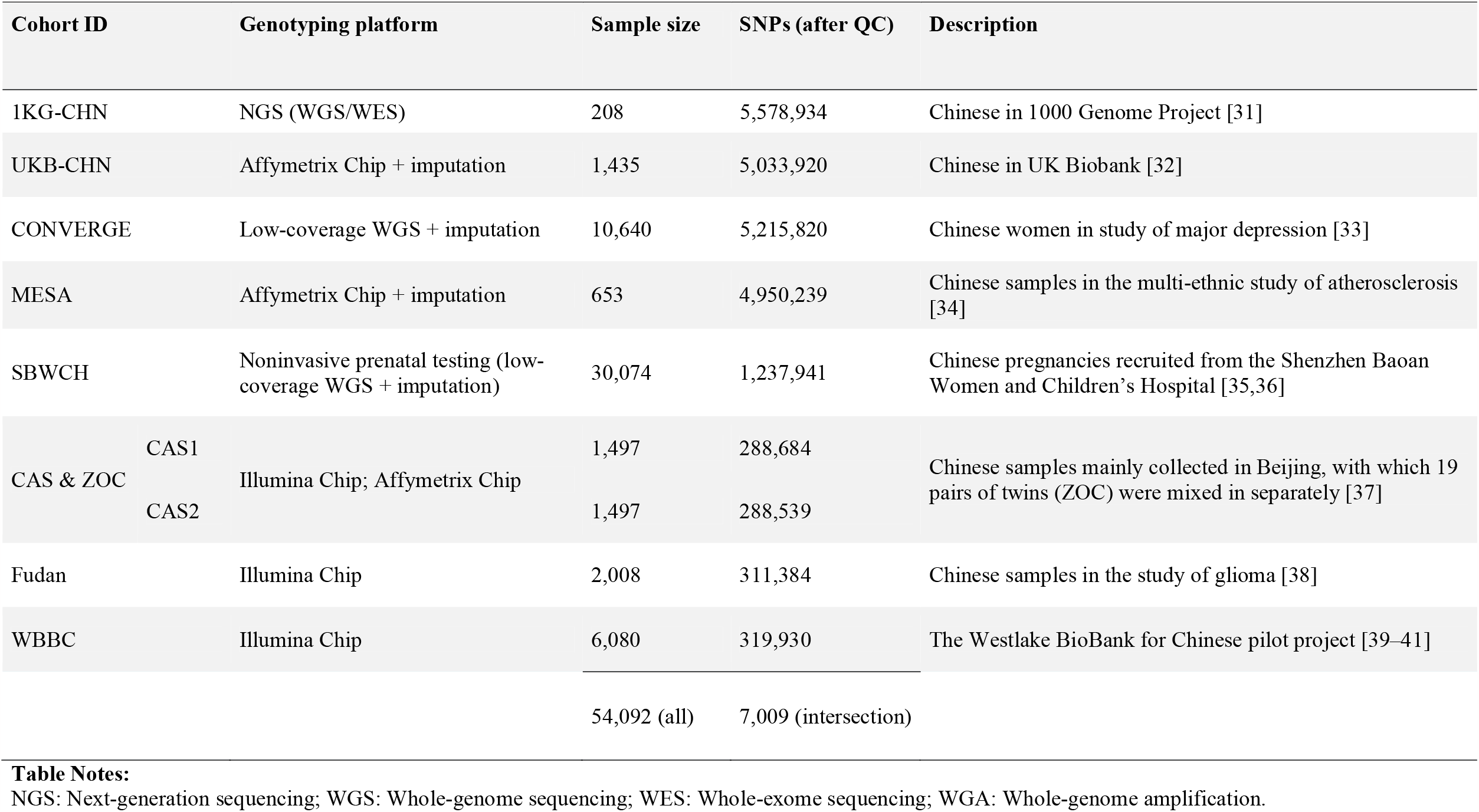
Summary information for the cohorts participated in this study.

#### 1KG-CHN (public)

We considered two Chinese subpopulations in 1000 Genome Project (1KG) [31], CHB (Han Chinese in Beijing, 103 individuals) and CHS (Southern Han Chinese, 105 individuals) as the reference population and also as a positive control in the cross-cohort test in Chinese datasets. Individuals in the project were genotyped by either whole-genome sequencing or whole-exome sequencing platform.

#### UKB-CHN (UKB application 41376)

The UKB includes 1,653 individuals of self-reported Chinese [32]. After genomic assessment, 1,435 were considered as Chinese origin. Individuals in the project were genotyped using the Applied Biosystems UK BiLEVE Axiom Array by Affymetrix, followed by the genotype imputation.

#### CONVERGE (public)

The CONVERGE consortium aimed to investigate major depressive disorder (MDD) [33]. It included 5,303 Chinese women with recurrent MDD and 5,337 controls, who were genotyped with low-coverage whole-genome sequencing followed by imputation.

#### MESA (accessible after dbGAP application)

The Multi-Ethnic Study of Atherosclerosis (MESA), which investigates subclinical cardiovascular disease [34], includes 653 Chinese samples, who were genotyped using Affymetrix Genome-Wide Human Single Nucleotide Polymorphism array 6.0, followed by genotype imputation.

#### SBWCH Biobank

The Shenzhen Baoan Women’s and Children’s Hospital (Baoan district, Shenzhen, Guangdong province) Biobank aims to investigate traits and diseases during pregnancy and at birth. 30,074 women were included in this study. Maternal genotypes were inferred from the non-invasive prenatal testing (NIPT) low depth whole genome sequencing data using STITCH [36] following the methodological pipeline that we previously published [35]. The average genotype imputation accuracy reaches 0.89 after filtration of INFO score 0.4.

#### CAS and ZOC

The Chinese Academy of Sciences (CAS) cohort is a prospective cohort study aiming to identify risk factors influencing physical and mental health of Chinese mental workers via a multi-omics approach. Since 2015, the study has recruited 4,109 CAS employees (48.2% male) located in Beijing, China. All participants belong to the research/education sector, and are characterized by a primary of Chinese Han origin (94.1%). DNA was extracted from peripheral blood samples and genotyped on the Infinium Asian Screening Array + MultiDisease-24 (ASA+MD) BeadChip, a specially designed genotyping array for clinical research of East Asian population with 743,722 variants. For validation purpose, samples were randomly split into CAS1 and CAS2. According to their records, ZOC was consisted of 19 homozygotic and heterozygotic siblings, who were evenly split into CAS1 and CAS2 as internal validation of encG-reg. ZOC is part of The Guangzhou Twin Eye Study (GTES), a prospective cohort study that included monozygotic and dizygotic twins born between 1987 and 2000 as well as their biological parents in Guangzhou, China. Baseline examinations were conducted in 2006, and all participants were invited to attend annual follow-up examinations. Non-fasting peripheral venous blood was collected by a trained nurse at baseline for DNA extraction, and genotyping was performed using the Affymetrix axiom arrays (Affymetrix) at the State Key Laboratory of Ophthalmology at Zhongshan Ophthalmic Center (ZOC) [37]. CAS and ZOC cohorts were deeply collaborated for certain studies, and consequently merged to fit this study.

#### Fudan

A multistage GWAS of glioma were performed in the Han Chinese population, with a total of 3,097 glioma cases and 4,362 controls. All Chinese Han samples used in this study were obtained through collaboration with multiple hospitals (Southern population from Huashan Hospital, Nanjing 1st Hospital, Northern population from Tiantan Hospital and Tangdu Hospital). DNA samples were extracted from blood samples and were genotyped using Illumina Human OmniExpress v1 BeadChips [38]. 2,008 samples were included for this study.

#### WBBC

The Westlake BioBank for Chinese (WBBC) cohort is a population-based prospective study with its major purpose to better understand the effect of genetic and environmental factors on growth and development from youngster to elderly [39]. The mean age of the study samples were 18.6 years for males and 18.5 years for females, respectively. The Westlake BioBank WBBC pilot project have finished whole-genome sequencing (WGS) in 4,535 individuals and high-density genotyping in 5,841 individuals [40,41].

The 9 Chinese datasets were reorganized into 9 cohorts (1KG-CHN, UKB-CHN, CONVERGE, META, SBWCH, CAS1, CAS2, Fudan, and WBBC) and to test ***encG-reg*** in the real world. Within CAS1 and CAS2, relatedness if identified by ***encG-reg*** would be verified by CAS. As would have been found, among other pairs of cohorts, sporadic relatedness might occur.

### Performance of encG-reg in Chinese cohorts

As summarized in **Fig 2A**, the Chinese cohort study was swiftly organized and completed within about 7 weeks, showing that ***encG-reg*** was an effective strategy with better ethical assurance. Following intra-cohort QCs and upon received summary information, we examined sample sizes and SNPs in each cohort (**Table 3**). In total, it included 54,092 samples and generated about 1 billion (N =930,140,004) pairs of tests. When allele frequencies were compared with that in CONVERGE, the majority of SNPs show consistent allele frequencies across cohorts (**S3 Fig and S6 Table**). The missing rates and the intersected SNPs were also examined across cohorts (**Figs S4-S5 and S7 Table**), after which a total of 7,009 SNPs were in common among 9 cohorts for the parsimony design of ***encG-reg*** (**Fig 6A**).

**Fig 6.**
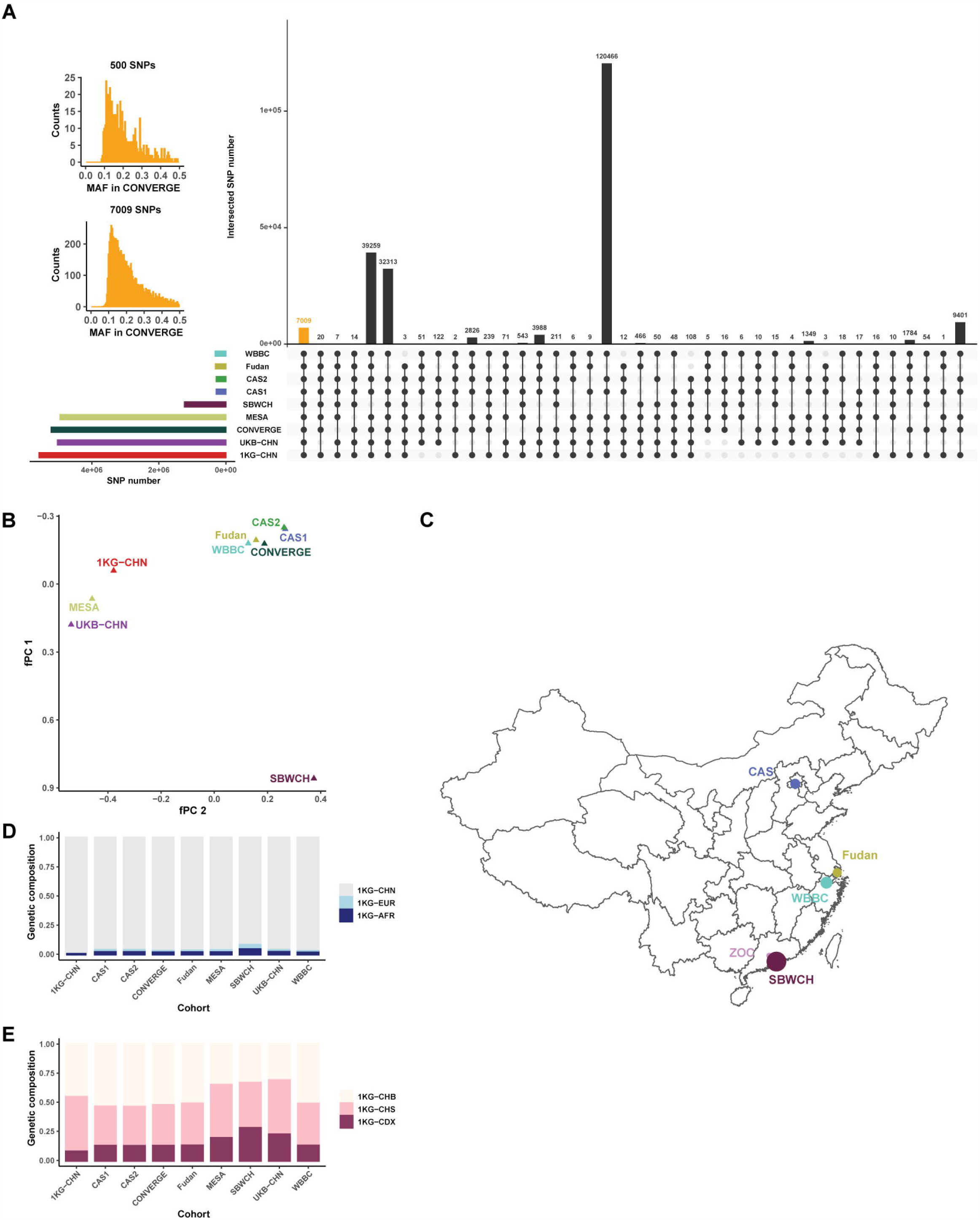
Cohort-level genetic background analyses for Chinese cohorts under parsimony encG-reg analysis. (**A**) Overview of the intersected SNPs across cohorts, a black dot indicated its corresponding cohort was included. Each row represented one cohort while each column represented one combination of cohorts. Dots linked by lines suggested cohorts in this combination. The height of bars represented the cohort’s SNP numbers (rows) or SNP intersection numbers (columns). Inset histogram plots show the distribution of the 7,009 intersected SNPs and the 500 SNPs randomly chosen from the 7,009 SNPs for ***encG-reg*** analysis. (**B**) 7,009 SNPs were used to estimate fPC from the intersection of SNPs for the 9 cohorts. Each triangle represented one Chinese cohort and was placed according to their first two principal component scores (fPC1 and fPC2) derived from the received allele frequencies. (**C**) Five private datasets have been pinned onto the base map from GADM (https://gadm.org/data.html) using R language. The size of point indicates the sample size of each dataset. (**D)** Global fStructure plot indicates global-level *F*_*st*_-derived genetic composite projected onto the three external reference populations: 1KG-CHN (CHB and CHS), 1KG-EUR (CEU and TSI), and 1KG-AFR (YRI), respectively; 4,296 of the 7,009 SNPs intersected with the three reference populations were used. (**E)** Within Chinese fStructure plot indicates within-China genetic composite. The three external references are 1KG-CHB (North Chinese), 1KG-CHS (South Chinese), and 1KG-CDX (Southwest minority Chinese Dai), respectively; 4,809 of the 7,009 SNPs intersected with these three reference populations were used. Along x axis are 9 Chinese cohorts and the height of each bar represents its proportional genetic composition of the three reference populations. Cohort codes: YRI, Yoruba in Ibadan representing African samples; CHB, Han Chinese in Beijing; CHS, Southern Han Chinese; CHN, CHB and CHS together; CEU, Utah Residents with Northern and Western European Ancestry; TSI, Tuscani in Italy; CDX, Chinese Dai in Xishuangbanna.

The results of fPCA and fStucture matched with their expected “geo-geno” mirror in Chinese samples [35]. The first eigenvector of fPCA distinguished southern and northern Chinese samples in this study: the SBWCH Biobank (dominantly sampled from Shenzhen, the southmost metropolitan city in mainland China) and CAS cohort (dominantly sampled from Beijing) (**Fig 6B and 6C**). Using a slightly different illustration strategy, the fStructure results, a counterpart to the well-known Structure plot in population genetics, were also consistent with the reported Chinese background of the 9 cohorts (**Fig 6D and 6E**).

We offered a list of 500 SNPs to be shared by the collaborators and estimated m_e_ to be 477 (evaluated in 1KG-CHN). The minimum number of k was 710, given the experiment-wise Type I error rate of 0.05 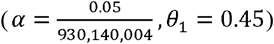 and the statistical power of 0.9. Each collaborator then encrypted their genotype matrix by the random matrix S. As foolproof controls, 1KG-CHN samples were consistently identified as “identical” inter-cohort.

Relatives were identified between CAS1 and CAS2, and SBWCH and WBBC (**Fig 7A** and **7B**). The pair-wise ***encG-reg*** distributions between cohorts were consistent with our theoretical expectation (**Fig 7C** and **S6**). For anticipated relatives, as each of the 19 Guangzhou twins was split into CAS1 and CAS2, 18 pairs were identified as monozygotic (MZ) or dizygotic (DZ) by ***encG-reg*** and verified by intra-cohort IBD calculation in CAS Beijing team (**Fig 7D)**. Remarkably, the pair of recorded twins that was not identified by ***encG-reg*** was verified as unrelated by IBD calculation, and ZOC team is conducting further investigation on potential logistic errors. These results demonstrated that ***encG-reg*** was reliable with well-controlled Type I and Type II error rates.

**Fig 7.**
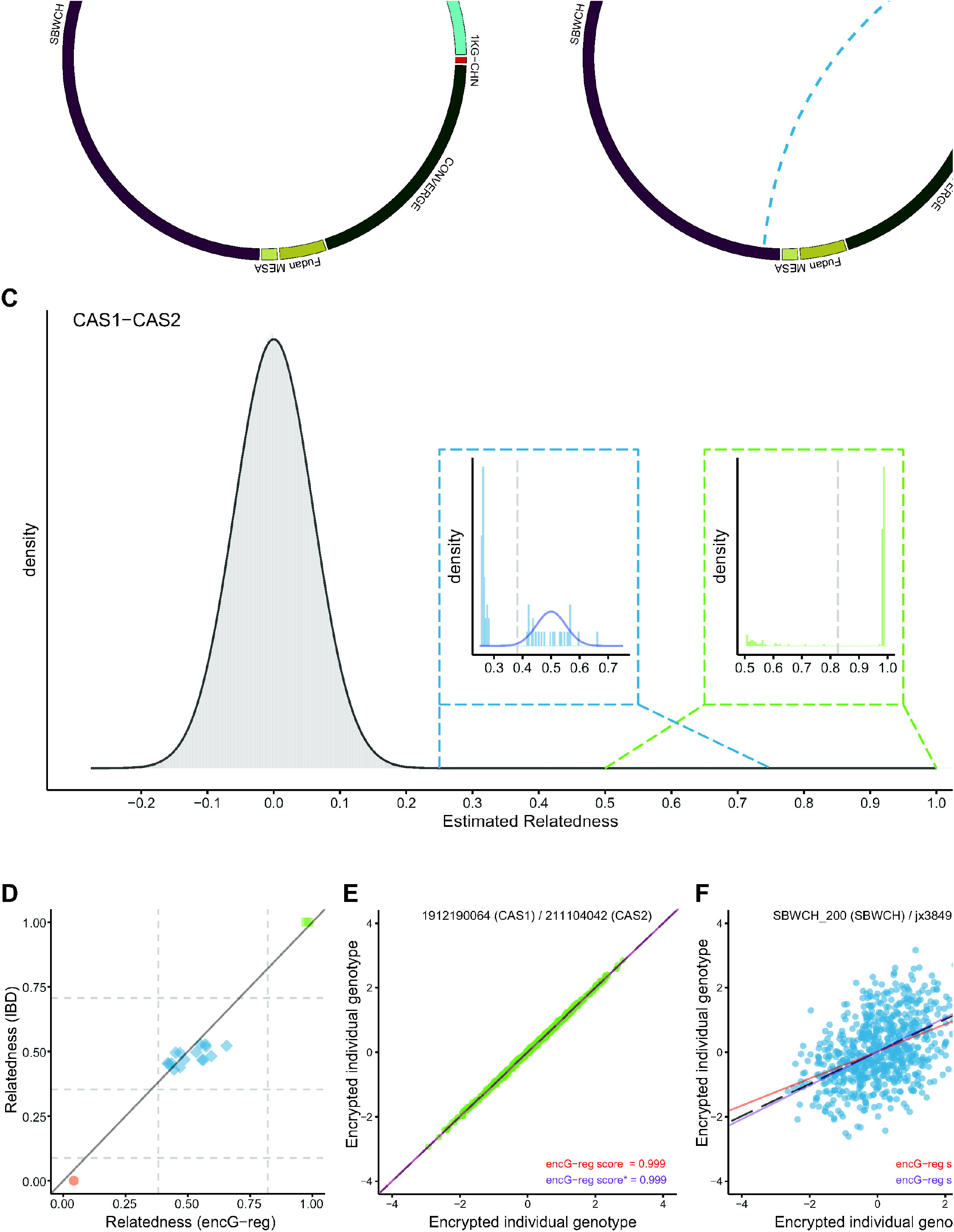
Detected identical pairs and 1st-degree pairs between Chinese cohorts. (**A**) The circle plot illustrates identical pairs and (**B**) 1st-degree pairs across 9 Chinses cohorts. The solid links indicates anticipated relatedness between the CAS cohorts. The dashed link indicates relatedness identified across cohorts. The length of each cohort bar is proportional to their respective sample sizes. (**C**) The histogram shows all estimated relatedness using ***encG-reg*** between CAS1 and CAS2, most of which are unrelated pairs and the theoretical probability density function is given as the normal distribution 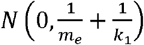 (grey solid curve). The inset histogram on the left shows the estimated relatedness around 0.5 and the theoretical probability density function is given as the normal distribution 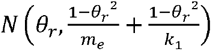 (blue solid curve). The threshold (grey dot line) for rejecting *H*_0_ was calculated by 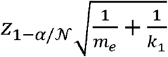. The inset histogram on the right shows the estimated relatedness around 1. The threshold (grey dot line) for rejecting *H*_0_ was calculated by 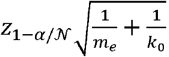. Here we included 208 controls merged from 1KG-CHN *m*_*e*_ = 477, *k*_1_ = 710, 𝒩 = 930,140,004. (**D**) Relationship verification for 19 Guangdong twins split in CAS cohorts. Dashed lines indicate inference criteria for detecting relatedness of different degrees. Solid line of y=x indicates the agreement between ***encG-reg*** and IBD. Points are colored with IBD-inferred relatedness in KING (identical in green, 1st-degree in blue, and unrelated in red) and are shaped according to ***encG-reg***-inferred relatedness (identical in square, 1st-degree in diamond, and unrelated in circle). (**E**) and (**F**) Illustration for ***encG-reg*** estimation for sporadic related inter-cohort samples. The grey line is the criterion for identical pairs (slope of 1) or 1st-degree pairs (slope of 0.5). The solid lines colored in red are without adjustment for missing values (***encG-reg*** score), and in the bottom (colored in purple) are adjusted for missing values (***encG-reg*** score^*^).

Particularly, we illustrated how sporadically related pairs were captured by ***encG-reg***. We detected 14 pairs of inter-cohort relatedness, including 4 pairs of identical samples and 10 pairs of 1st-degree relatives (**Table 4**). For these sporadic related inter-cohort samples, ***encG-reg*** exhibited their relatedness in the forms of regression plots and estimated regression coefficients, two examples were given in **Fig 7E** and **7F**. Thirteen out of fourteen pairs of sporadic related pairs were identified between CAS1 and CAS2. This could likely be attributed to the centralized sampling of the CAS cohort among CAS employees, as it is highly probable that family members are intended to work in the same company. Moreover, despite the higher missing rate in SBWCH, which introduced additional noise, ***encG-reg*** still managed to capture one across-cohort relatives between SBWCH and WBBC. To avoid possible breaching of privacy, we refrained from further exploring their relationship as extensively as we did for UKB.

**Table 4.** Supporting evidence for sporadic related pairs.

| Pair | Cohort 1 | ID 1 | Cohort 2 | ID 2 | Score (SD<sup>a</sup>) | Adjusted Score<sup>b</sup> (SD) | Inferred relatedness |
| --- | --- | --- | --- | --- | --- | --- | --- |
| 1 | CAS1 | 1912190064 | CAS2 | 211104042 | 0.999 (0.001) | 0.999 (0.001) | Identity |
| 2 | CAS1 | 2106190041 | CAS2 | 2009111151 | 0.999 (0.002) | 0.999 (0.002) | Identity |
| 3 | CAS1 | 211119018 | CAS2 | 2010130356 | 0.999 (0.001) | 0.999 (0.001) | Identity |
| 4 | CAS1 | 20091209 | CAS2 | 2011050139 | 0.999 (0.001) | 0.999 (0.001) | Identity |
| 5 | CAS1 | 20090801 | CAS2 | 21110202 | 0.421 (0.034) | 0.421 (0.034) | 1st-degree |
| 6 | CAS1 | 20090801 | CAS2 | 2016122301 | 0.421 (0.034) | 0.421 (0.034) | 1st-degree |
| 7 | CAS1 | 211029082 | CAS2 | 211029076 | 0.792 (0.023) | 0.792 (0.023) | 1st-degree |
| 8 | CAS1 | 1912160050 | CAS2 | 211104131 | 0.561 (0.031) | 0.561 (0.031) | 1st-degree |
| 9 | CAS1 | 211104139 | CAS2 | 211104138 | 0.458 (0.033) | 0.458 (0.033) | 1st-degree |
| 10 | CAS1 | 211104147 | CAS2 | 211104148 | 0.513 (0.032) | 0.513 (0.032) | 1st-degree |
| 11 | CAS1 | 211104174 | CAS2 | 211104173 | 0.531 (0.032) | 0.531 (0.032) | 1st-degree |
| 12 | CAS1 | 211104198 | CAS2 | 211104199 | 0.508 (0.032) | 0.508 (0.032) | 1st-degree |
| 13 | CAS1 | 211104164 | CAS2 | 211104236 | 0.415 (0.034) | 0.415 (0.034) | 1st-degree |
| 14 | SBWCH | SBWCH_200 | WBBC | jx3849 | 0.420 (0.034) | 0.524 (0.043) | 1st-degree |

**Table Notes:** IDs were de-identified by each cohort.
<sup>a</sup>Standard deviation (SD) is calculated from $SD_{b_{ij}} = \sqrt{\frac{cov(\hat{\mathbf{x}}_i, \hat{\mathbf{x}}_j)}{var(\hat{\mathbf{x}}_i)}}$ , where $\hat{\mathbf{x}}_i$ and $\hat{\mathbf{x}}_j$ are the vectors of the encrypted genotypes for two individuals.
<sup>b</sup>Due to missing data, the corrected score, is adjusted for the genotype missing rate between the pair of individuals

## Discussion

One of the early attempts on detecting cross-cohort relatives was limited to detecting overlapping individuals by one-way cryptographic hashes, which offered qualitative but not quantitative conclusions on false positive and false negative rates [13]. To settle the question of exact encryption precision, we focused on investigating the intrinsic consequence after genotype encryption with a random matrix and proposed ***encG-reg***. We described the properties of ***encG-reg*** in how *k* and *m*_*e*_ influence its precision. This property was well testified in both the UKB example and the collaboration across China. Our investigation led to controllable encryption precision even under a variety of genotype platforms and datasets with different qualities. It should be noticed, as a proof-of-concept, we only studied additive GRM, which corresponds to *IBD*_1_. Obviously, our work can be extended to dominance-GRM (or two-allele “IBD=2” scheme), so as to further split 1st-degree relatives into parent-offspring (*IBD*_0_ = 0, *IBD*_1_= 1.0, and *IBD*_2_ = 0) and full sibs (*IBD*_0_ = 0.25, *IBD*_1_ = 0.5, and *IBD*_2_ = 0.25). To this point, we have only presented the outcomes concerning identity, 1st-degree, and 2nd-degree relatedness in UKB. This is primarily due to the absence of a distinct definition for true relatedness. To conduct further examination for the exact inference of distant relatives, a dataset with more pedigree information should be employed and a study design with more comprehensive comparisons should be considered [42].

As demonstrated in UKB multi-ethnicity samples, ***encG-reg*** could be applied for biobank-scale datasets with high precision in comparing with conventional individual-level benchmark methods such as KING and GRM. The evaluation using Chinese cohorts is the first attempt to develop an encrypted method that can be applied in large-scale searching relatives with encrypted genomic data. In this experiment, for convenience and manageability, we only considered parsimony design by using shared SNPs across the 9 Chinese cohorts. Switching to exhaustive design will be a better option if each pair of cohorts conducts ***encG-reg*** for their customized degree of relatives.

For either exhaustive design or parsimony design of ***encG-reg***, the core algorithm is algebraic and requires little human information in its implementation. Thus, an automatic central analysis facility that can significantly host and synchronize more cohorts will be attractive. An exhaustive implementation of ***encG-reg*** will search even deeper relatedness across cohorts in a highly mobilizing nation like China, in which relatives used to live nearby but now are distant due to industrialization [43]. A much deeper implementation of ***encG-reg*** will bring out unique resource for conducting biomedical research at large scale as including familial information as demonstrated [44]. Last but not least, ***encG-reg*** is a developed tool that is with much better protected genomic privacy, and can facilitate necessary relative searching when it is needed. It is not purposed to penetrate membership or other unethical activities.

An attack on the central site may result in the leak of encrypted genotype matrices and estimated relatedness score, but no raw genotype matrices will be leaked. However, since the individual IDs were de-identified by each cohort, such as individual IDs presented in **Table 4**, no more information can be traced back by other sites or the central site. Moreover, additional secure protection can be implemented at the central site (which can be a cloud server), and this is about the design on an entrusted server. In the worst-case scenario—a colluded central analyst, both **S** and **XS** have been leaked, certain adversary attacks, such as PCA-attack, may be carried out. In PCA-attack, **X** can be adversary reconstructed via principal component analysis on **XS**, if the correlation structure of SNPs can be approximated in a proper reference population [45]. To mitigate the risk of a PCA-attack, one straightforward defence is to use SNPs that are in linkage equilibrium as much as possible (*m* ≈ *m*_*e*_ then), and it ensures that the correlation matrix closely approximates the identity matrix. As noticed, the strategy of masking the original genotype after multiplication of **S** resembles matrix random projection employed in classifying the transactions as legitimate or fraudulent across financial institutions, and a class of methods of possibly reconstructing **X** have been discussed [46]. Consequently, we are confident for the safety of the whole practice, for now and for the future when the ***encG-reg*** grows to a more broadly utilized application.

The homomorphic encryption such as CKKS scheme [47] and Fan and Vercauteren (FV) scheme [48] have recently been employed in developing HE-KING [49,50], which are also applicable to our study. Nevertheless, the computational cost of HE-KING is substantial, for which after encryption the memory cost is often one or two orders of magnitude of the original genotypes. **Eq 3** provides a lower bound of SNP numbers for detecting relatedness and consequently can be plugged into HE-KING, a useful quantity that is able to reduce the SNP number and computational cost in particular when analyzing biobank-scale data such as UKB.

## Supporting information

SuppNoteFigsTables

## Acknowledgements

We thank the participants of the included cohorts and of UK Biobank for making this work possible (UKB application 41376). Thank Miss Qiu Feng, Mr Mu Wentao, Dr Qian Jie, and Mr Mei Lixiao for various assistance in making this work possible. Thanks Professor Xiaoqian Jiang at the University of Texas Health Science Center at Houston for helpful discussion on HE-KING. Our reviewers brought to our attention the random projection technique, which has consequently prompted us to consider the potential for various adversary attacks.

## Supporting Information captions

**S1 Text. Details about the derivations of sampling variance, details about the selection of ethnicity-insensitive SNPs, and details about fPCA and fStructure**.

This appendix contains supplementary notes on “Details about random matrix properties for encGRM” (Note 1), “The variance of GRM (assumption: multivariate normal distribution)” (Note 2), “The variance of GRM (assumption: binomial distribution)” (Note 3), “The variance of encGRM” (Note 4), “The variance of ***encG-reg***” (Note 5), “Details about power calculation in Eq 4” (Note 6), “Selection of ethnicity-insensitive SNPs” (Note 7), and “fPCA and fStructure” (Note 8).

**S1 Fig. Heatmap presenting the role random matrix played in matrix multiplication**.

We generate a random matrix **S** _*m* × *k*_ sampling from *N*(0,1/*k*) and plot **SS**^T^ (**B**) against an identity matrix (**A**). We also generate two small populations containing 20 and 25 individuals, respectively. Their genotype matrices are noted as **X**_1_ and **X**_2_, and plot the matrix multiplication product 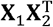 before (**C**) and after encryption 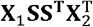 (**D**). The column number for the random matrix is *k*=500 and the number of SNPs is *m*=100.

**S2 Fig. Validation for the sampling variance of GRM (assumption: binomial distribution)**.

To testify the variance of GRM under the assumption of binomial distribution, we simulated 1,000 pairs of different degrees of relatives, and 2,000 markers with same MAF from 0.05 to 0.45 per increase in 0.1. We compared the observed variance of relatedness with the theoretical relatedness in 10 repeats.

**S3 Fig. Comparison of MAF in each cohort with the reference panel CONVERGE**.

Comparison of MAF in CONVERGE with the frequency of the same allele in each cohort. Each hexagonal bin is colored according to the number of markers falling in that bin.

**S4 Fig. Comparison of MAF in CONVERGE and in each cohort (7**,**009 shared SNPs)**.

Comparison of MAF in CONVERGE with the frequency of the same allele in each Chinese cohort, considering 7,009 overlapping SNPs only. Each hexagonal bin is colored according to the number of markers falling in that bin.

**S5 Fig. Non-missing allele counts in Chinese cohorts**.

Distributions of non-missing allele counts in each cohort. Maximum allele counts = 2*m* = 1,000.

**S6 Fig. Distribution of encG-reg score across Chinese cohorts**.

The histogram shows all estimated relatedness using ***encG-reg*** between SBWCH and WBBC, most of which are unrelated pairs and the theoretical probability density function is given as the normal distribution 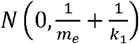 (grey solid curve). The inset histogram on the left shows the estimated relatedness around 0.5 and the theoretical probability density function is given as the normal distribution 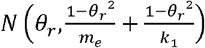 (blue solid curve). The threshold (grey dot line) for rejecting *H*_0_ was calculated by 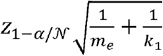. The inset histogram on the right shows the estimated relatedness around 1. The threshold (grey dot line) for rejecting *H*_0_ was calculated by 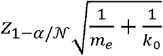. Here we included 208 controls merged from 1KG-CHN. *m*_*e*_ = 477, *k*_0_ = 70, *k*_1_ = 710, 𝒩 = 930,140,004.

**S1 Table. Notation definitions**.

**S2 Table**. *m*_*e*_ **estimation in UK Biobank and Chinese cohorts**.

**S3 Table. Sample size for 19 cohorts in UKB**.

**S4 Table. The number of ethnicity-insensitive SNPs intersected between each pair of UKB cohorts and the elapsed time when performing encG-reg**.

**S5 Table. The number of inferred relatedness at exhaustive and parsimony designs. S6 Table. Summary of cross-cohort quality control**.

**S7 Table. The number of intersected SNPs between each pair of Chinese cohorts**.

