## Supplementary material for "Searching across-cohort relatives in 54,092 GWAS samples via encrypted genotype regression": SuppNoteFigsTables: S1_Table.docx

**S1 Table. Notation definitions.**

| **Notation** | **Definition** |
| --- | --- |
| $n_{1}$ | Sample size of cohort 1. |
| $n_{2}$ | Sample size of cohort 2. |
| $\mathbf{X}_{1}$ | The scaled additive (coded as 0, 1, or 2 defining the number of reference alleles) genotype matrix of cohort 1. |
| $\mathbf{X}_{2}$ | The scaled additive genotype matrix of cohort 2. |
| $\mathbf{S}$ | A matrix of $m$ rows and $k$ columns, each element is sampled from $N\left( 0,\sigma^{2} \right)$. |
| ${\hat{\mathbf{X}}}_{1}$ | The encrypted additive genotype matrix of cohort 1, ${\hat{\mathbf{X}}}_{1}=\mathbf{X}_{1}\mathbf{S}$. |
| ${\hat{\mathbf{X}}}_{2}$ | The encrypted additive genotype matrix of cohort 2, ${\hat{\mathbf{X}}}_{2}=\mathbf{X}_{2}\mathbf{S}$. |
| $m$ | The number of markers, also represents the number of rows for random matrix $\mathbf{S}$. |
| $m_{e}$ | The number of effective markers, which has accounted for LD between markers. |
| $k$ | The iteration number for randomization, as represented the number of columns for random matrix $\mathbf{S}$. |
