## Supplementary material for "Searching across-cohort relatives in 54,092 GWAS samples via encrypted genotype regression": SuppNoteFigsTables: S1_Text_Nov10.docx

### Supplementary Notes

#### Note 1 Details about random matrix properties for encGRM

Prove: when $\mathbf{S}$ is an $m$-by-$k$ matrix whose elements are independently sampled from a normal distribution $s_{ij}\sim N\left( 0,\sigma^{2} \right)$, then

$$E\left( \mathbf{S}\mathbf{S}^{T} \right)=k\sigma^{2}\mathbf{I}$$

Proof:

The expectation of element on the diagonal of $\mathbf{S}\mathbf{S}^{T}$ is,

$$E\left( \sum_{j}^{k} s_{ij}^{2} \right)=\sum_{j}^{k} E(s_{ij}^{2})=k\sigma^{2}$$

The expectation of element of the $i^{th}$ row and $j^{th}$ column of $\mathbf{S}\mathbf{S}^{T}$ is,

$$E\left( \sum_{j}^{k} s_{ij}s_{i^{'}j} \right)=\sum_{j}^{k} E(s_{ij}s_{i^{'}j})=0$$

#### Note 2 The variance of GRM (assumption: multivariate normal distribution)

We assume the genotypes are scaled by their frequencies to zero mean and a unit variance, $E\left( x_{im} \right)=0$ and $var\left( x_{im} \right)=1$. The correlation between $x_{im}$ and $x_{jm}$ is $\theta$, or equivalently considered as identical by descent. Therefore, $E\left( x_{im}x_{jm} \right)=cov\left( x_{im},x_{jm} \right)=cor(x_{im},x_{jm})=\theta$.

The genetic relatedness score is defined as,

$$g_{ij}=\frac{1}{M}\sum_{m=1}^{M} x_{im}x_{jm}$$

**Expectation**:

$$E\left( g_{ij} \right)=\frac{1}{M}E\left( \sum_{m=1}^{M} x_{im}x_{jm} \right)=\frac{1}{M}\sum_{m=1}^{M} E\left( x_{im}x_{jm} \right)=\theta$$

**Variance**:

$$E\left( g_{ij}^{2} \right)=\frac{1}{M^{2}}E\left[ \left( \sum_{m=1}^{M} x_{im}x_{jm} \right)^{2} \right]=\frac{1}{M^{2}}\left[ \sum_{m=1}^{M} E\left( x_{im}^{2}x_{jm}^{2} \right)+\sum_{m}^{M} \sum_{m^{'}\neq m}^{M} E(x_{im}x_{jm}x_{im^{'}}x_{jm^{'}}) \right]$$

Here, according to Isserlis’s theorem,

$$E\left( x_{im}^{2}x_{jm}^{2} \right)=E\left( x_{im}^{2} \right)E\left( x_{jm}^{2} \right)+2E^{2}\left( x_{im}x_{jm} \right)=1+2\theta^{2}$$

$$E\left( x_{im}x_{jm}x_{im^{'}}x_{jm^{'}} \right)=E\left( x_{im}x_{jm} \right)E\left( x_{im^{'}}x_{jm^{'}} \right)=\theta^{2}$$

Therefore, the variance of GRM is,

$$var\left( g_{ij} \right)=E\left( g_{ij}^{2} \right)-E^{2}\left( g_{ij} \right)=\frac{1}{M^{2}}\left[ M\left( 1+2\theta^{2} \right)+M\left( M-1 \right)\theta^{2} \right]-\theta^{2}=\frac{1+\theta^{2}}{M}$$

When considering LD between SNPs,

$$E\left( x_{im}x_{jm}x_{im^{'}}x_{jm^{'}} \right)=E\left( x_{im}x_{jm} \right)E\left( x_{im^{'}}x_{jm^{'}} \right)+E\left( x_{im}x_{im^{'}} \right)E\left( x_{jm}x_{jm^{'}} \right)+E\left( x_{im}x_{jm^{'}} \right)E\left( x_{jm}x_{im^{'}} \right)$$

$$=\theta^{2}+\rho_{mm^{'}}^{2}+\theta^{2}\rho_{mm^{'}}^{2}$$

To note that the effective number of markers is defined as, $\frac{1}{M_{e}}=\frac{\sum_{m}^{M} \sum_{m^{'}}^{M} \rho_{mm^{'}}^{2}}{M^{2}}=\frac{M+\sum_{m}^{M} \sum_{m^{'}\neq m}^{M} \rho_{mm^{'}}^{2}}{M^{2}}$. Therefore, the variance of GRM becomes,

$$var\left( g_{ij} \right)=E\left( g_{ij}^{2} \right)-E^{2}\left( g_{ij} \right)$$

$$=\frac{1}{M^{2}}\left[ M\left( 1+2\theta^{2} \right)+\sum_{m}^{M} \sum_{m^{'}\neq m}^{M} \left( \theta^{2}+\rho_{mm^{'}}^{2}+\theta^{2}\rho_{mm^{'}}^{2} \right) \right]-\theta^{2}$$

$$=\frac{1}{M^{2}}\left[ M\left( 1+2\theta^{2} \right)+M\left( M-1 \right)\theta^{2}+\left( 1+\theta^{2} \right)\sum_{m}^{M} \sum_{m^{'}\neq m}^{M} \rho_{mm^{'}}^{2}-M^{2}\theta^{2} \right]$$

$$=\frac{1}{M^{2}}\left[ M+M\theta^{2}+\left( 1+\theta^{2} \right)\left( \frac{M^{2}}{M_{e}}-M \right) \right]$$

$$=\frac{1+\theta^{2}}{M_{e}}$$

#### Note 3 The variance of GRM (assumption: binomial distribution)

Although the variance of GRM can be derived under the assumption of multivariate normal distribution in the last section, it needs to be modeled at binomial distribution if we take the allele frequency into account. If we consider two diploid individuals, each with two alleles ($A$ and $a$) at one locus. The allele frequencies are $p$ and $q$ for allele $A$ and $a$, respectively. Let $\theta$ denote the probability that two alleles sampled at random are identical by descent (IBD). For a pair of relatives, the probabilities of two alleles conditioning on the probability of IBD $\theta$ is:

| **Allelic pair for relatives** | **Probability** |
| --- | --- |
| $A/A$ | $\theta p+\left( 1-\theta\right)p^{2}$ |
| $A/a$ | $\left( 1-\theta\right)pq$ |
| $a/A$ | $\left( 1-\theta\right)pq$ |
| $a/a$ | $\theta q+\left( 1-\theta\right)q^{2}$ |

The above table is applied to their respect second allele pair. The probabilities of sixteen genotype combinations between two individuals are filled in this $4\times4$ table, which is symmetric:

|  |  | First allelic pair | | | |
| --- | --- | --- | --- | --- | --- |
|  | **Probability** | $A/A$ | $A/a$ | $a/A$ | $a/a$ |
| Second allelic  pair | $A/A$ | $\left[ \theta p+\left( 1-\theta\right)p^{2} \right]^{2}$ | $\left( 1-\theta\right)pq$  $[\theta p+\left( 1-\theta\right)p^{2}]$ | $\left( 1-\theta\right)pq$  $[\theta p+\left( 1-\theta\right)p^{2}]$ | $\left[ \theta p+\left( 1-\theta\right)p^{2} \right]$  $[\theta q+\left( 1-\theta\right)q^{2}]$ |
|  | $A/a$ |  | $\left( 1-\theta\right)^{2}p^{2}q^{2}$ | $\left( 1-\theta\right)^{2}p^{2}q^{2}$ | $\left( 1-\theta\right)pq$  $[\theta q+\left( 1-\theta\right)q^{2}]$ |
|  | $a/A$ |  |  | $\left( 1-\theta\right)^{2}p^{2}q^{2}$ | $\left( 1-\theta\right)pq$  $[\theta q+\left( 1-\theta\right)q^{2}]$ |
|  | $a/a$ | Symmetric |  |  | $\left[ \theta q+\left( 1-\theta\right)q^{2} \right]^{2}$ |

Ignoring the order of two individuals, we can recalculate the frequencies by summing up the corresponding probabilities in each cell. As individual genotypes are standardized by $\hat{x}=\frac{x-2p}{\sqrt{2pq}}$, where $x$ is coded as the number of minor alleles. Relatedness score between individual $i$ and individual $j$ is estimated by $\frac{(x_{i}-2\hat{p})(x_{j}-2\hat{p})}{2\hat{p}\hat{q}}$. $\hat{p}$ and $\hat{q}$ are the estimated allele frequencies of $A$ and $a$. The rearranged probabilities and relatedness scores are:

| **Genotype pair** | **Genotype code** | **Probability (**$\boldsymbol{f}_{\boldsymbol{i}}$**)** | **Relatedness (**$\boldsymbol{X}_{\boldsymbol{i}}$**)** |
| --- | --- | --- | --- |
| $\{AA, AA\}$ | $\{2, 2\}$ | $\left[ \theta p+\left( 1-\theta\right)p^{2} \right]^{2}$ | $\frac{\left( 2-2p \right)^{2}}{2pq}$ |
| $\{AA, Aa\}$ | $\{2, 1\}$ | $4\theta\left( 1-\theta\right)p^{2}q+4\left( 1-\theta\right)^{2}p^{3}q$ | $\frac{\left( 2-2p \right)\left( 1-2p \right)}{2pq}$ |
| $\{AA, aa\}$ | $\{2, 0\}$ | $2\left( 1-\theta\right)^{2}p^{2}q^{2}$ | $-\frac{2p\left( 2-2p \right)}{2pq}$ |
| $\{Aa, Aa\}$ | $\{1, 1\}$ | $4\left( 1-\theta\right)^{2}p^{2}q^{2}+2\theta pq$ | $\frac{\left( 1-2p \right)^{2}}{2pq}$ |
| $\{Aa, aa\}$ | $\{1, 0\}$ | $4\theta\left( 1-\theta\right)pq^{2}+4\left( 1-\theta\right)^{2}pq^{3}$ | $-\frac{2p\left( 1-2p \right)}{2pq}$ |
| $\{aa, aa\}$ | $\{0, 0\}$ | $\left[ \theta q+\left( 1-\theta\right)q^{2} \right]^{2}$ | $\frac{\left( -2p \right)^{2}}{2pq}$ |

$E\left( Relatedness \right)=\sum X_{i}f_{i}=\theta$ and $var\left( Relatedness \right)=\sum X_{i}^{2}f_{i}- \sum X_{i}f_{i}=1-2\theta+\frac{\theta}{2pq}$

If we consider $m$ loci of the same allele frequency, the variance of relatedness becomes $\frac{1-2\theta+\frac{\theta}{2pq}}{m}$. Therefore, the more $p$ approaches 0.5, the smaller variance of relatedness is.

Whether the sampling variance is derived from normal or binomial distribution, their results are similar. Simulation results see **Figure S3**.

#### Note 4 The variance of encGRM

After multiplying genotype matrix by a random matrix $\mathbf{S}=\left\{ s_{mk} \right\}\sim N(0,\frac{1}{M})$, the encrypted genotypes for individual $i$ and individual $j$ are $\hat{x}_{ik}=\sum_{m}^{M} x_{im}s_{mk}$ and $\hat{x}_{jk}=\sum_{m}^{M} x_{jm}s_{mk}$. The encrypted genetic relatedness score (encGRM) is defined as,

$$\hat{g}_{ij}=\frac{1}{K}\sum_{k=1}^{K} \hat{x}_{ik}\hat{x}_{jk}$$

**Expectation:**

$$E\left( \hat{g}_{ij} \right)=\frac{1}{K}E\left( \sum_{k}^{K} (\sum_{m}^{M} x_{im}s_{mk})(\sum_{m}^{M} x_{jm}s_{mk}) \right)=\frac{1}{K}\sum_{k}^{K} \sum_{m}^{M} \sum_{m^{'}}^{M} E\left( x_{im}s_{mk}x_{jm^{'}}s_{m^{'}k} \right)$$

When $m=m^{'}$, $E\left( x_{im}s_{mk}x_{jm^{'}}s_{m^{'}k} \right)=E\left( x_{im}x_{jm} \right)E\left( s_{mk}^{2} \right)=\theta/M$, and we have.

$$E\left( \hat{g}_{ij} \right)=\frac{1}{K}\times K\times M\times\frac{\theta}{M}=\theta$$

**Variance:**

$$E\left( \hat{g}_{ij}^{2} \right)=\frac{1}{K^{2}}E\left[ \left( \sum_{k}^{K} (\sum_{m}^{M} x_{im}s_{mk})(\sum_{m}^{M} x_{jm}s_{mk}) \right)^{2} \right]=\frac{1}{K^{2}}E\left[ \left( \sum_{k}^{K} \sum_{m}^{M} \sum_{m^{'}}^{M} x_{im}s_{mk}x_{jm^{'}}s_{m^{'}k} \right)^{2} \right]=\frac{1}{K^{2}}\sum_{k_{1}}^{K} \sum_{k_{2}}^{K} \sum_{m_{1}}^{M} \sum_{m_{2}}^{M} \sum_{m_{3}}^{M} \sum_{m_{4}}^{M} E\left( x_{im_{1}}s_{m_{1}k_{1}}x_{jm_{2}}s_{m_{2}k_{1}}x_{im_{3}}s_{m_{3}k_{2}}x_{jm_{4}}s_{m_{4}k_{2}} \right)$$

There are only two situations when the expectation of element is not equal to zero.

$E\left( x_{im_{1}}^{2} \right)E\left( x_{jm_{2}}^{2} \right)E\left( s_{m_{1}k_{1}}^{2} \right)E\left( s_{m_{2}k_{2}}^{2} \right)=\frac{1}{M^{2}}$ and $E\left( x_{im_{1}}x_{jm_{1}} \right)E\left( x_{im_{2}}x_{jm_{2}} \right)E\left( s_{m_{1}k_{1}}^{2} \right)E\left( s_{m_{2}k_{2}}^{2} \right)=\frac{\theta^{2}}{M^{2}}$

**I** when $m_{1}=m_{2}=m_{3}=m_{4}$

$$E\left( x_{im_{1}}s_{m_{1}k_{1}}x_{jm_{2}}s_{m_{2}k_{1}}x_{im_{3}}s_{m_{3}k_{2}}x_{jm_{4}}s_{m_{4}k_{2}} \right)=E\left( x_{im_{1}}^{2}x_{jm_{1}}^{2}s_{m_{1}k_{1}}^{2}s_{m_{1}k_{2}}^{2} \right)$$

**I-1** $k_{1}=k_{2}$:

$$E\left( x_{im_{1}}^{2}x_{jm_{1}}^{2}s_{m_{1}k_{1}}^{4} \right)=3E\left( x_{im_{1}}^{2} \right)E\left( x_{im_{1}}^{2} \right)E\left( s_{m_{1}k_{1}}^{2} \right)E\left( s_{m_{1}k_{1}}^{2} \right)+6E\left( x_{im_{1}}x_{jm_{1}} \right)E\left( x_{im_{1}}x_{jm_{1}} \right)E\left( s_{m_{1}k_{1}}^{2} \right)E\left( s_{m_{1}k_{1}}^{2} \right)$$

$$=\frac{3+6\theta^{2}}{M^{2}}$$

**I-2** $k_{1}\neq k_{2}$:

$$E\left( x_{im_{1}}^{2}x_{jm_{1}}^{2}s_{m_{1}k_{1}}^{2}s_{m_{1}k_{2}}^{2} \right)=E\left( x_{im_{1}}^{2} \right)E\left( x_{jm_{1}}^{2} \right)E\left( s_{m_{1}k_{1}}^{2} \right)E\left( s_{m_{1}k_{2}}^{2} \right)+2E\left( x_{im_{1}}x_{jm_{1}} \right)E\left( x_{im_{1}}x_{jm_{1}} \right)E\left( s_{m_{1}k_{1}}^{2} \right)E\left( s_{m_{1}k_{2}}^{2} \right)$$

$$=\frac{1+2\theta^{2}}{M^{2}}$$

**II** when $m_{1}=m_{2}, m_{3}=m_{4}$

$$E\left( x_{im_{1}}s_{m_{1}k_{1}}x_{jm_{2}}s_{m_{2}k_{1}}x_{im_{3}}s_{m_{3}k_{2}}x_{jm_{4}}s_{m_{4}k_{2}} \right)=E\left( x_{im_{1}}x_{im_{3}}x_{jm_{1}}x_{jm_{3}}s_{m_{1}k_{1}}^{2}s_{m_{3}k_{2}}^{2} \right)$$

**II-1** $k_{1}=k_{2}$:

$$E\left( x_{im_{1}}x_{im_{3}}x_{jm_{1}}x_{jm_{3}}s_{m_{1}k_{1}}^{2}s_{m_{3}k_{1}}^{2} \right)=E\left( x_{im_{1}}x_{jm_{1}} \right)E\left( x_{im_{3}}x_{jm_{3}} \right)E\left( s_{m_{1}k_{1}}^{2} \right)E\left( s_{m_{3}k_{1}}^{2} \right)=\frac{\theta^{2}}{M^{2}}$$

**II-2** $k_{1}\neq k_{2}$:

$$E\left( x_{im_{1}}x_{im_{3}}x_{jm_{1}}x_{jm_{3}}s_{m_{1}k_{1}}^{2}s_{m_{3}k_{2}}^{2} \right)=E\left( x_{im_{1}}x_{jm_{1}} \right)E\left( x_{im_{3}}x_{jm_{3}} \right)E\left( s_{m_{1}k_{1}}^{2} \right)E\left( s_{m_{3}k_{2}}^{2} \right)=\frac{\theta^{2}}{M^{2}}$$

**III** when $m_{1}=m_{3}, m_{2}=m_{4}$,

$$E\left( x_{im_{1}}s_{m_{1}k_{1}}x_{jm_{2}}s_{m_{2}k_{1}}x_{im_{3}}s_{m_{3}k_{2}}x_{jm_{4}}s_{m_{4}k_{2}} \right)=E\left( x_{im_{1}}^{2}x_{jm_{2}}^{2}s_{m_{1}k_{1}}s_{m_{2}k_{1}}s_{m_{1}k_{2}}s_{m_{2}k_{2}} \right)$$

**III-1** $k_{1}=k_{2}$:

$$E\left( x_{im_{1}}^{2}x_{jm_{1}}^{2}s_{m_{1}k_{1}}^{2}s_{m_{2}k_{1}}^{2} \right)=E\left( x_{im_{1}}^{2} \right)E\left( x_{jm_{1}}^{2} \right)E\left( s_{m_{1}k_{1}}^{2} \right)E\left( s_{m_{2}k_{1}}^{2} \right)=\frac{1}{M^{2}}$$

**IV** when $m_{1}=m_{4}, m_{2}=m_{3}$

$$E\left( x_{im_{1}}s_{m_{1}k_{1}}x_{jm_{2}}s_{m_{2}k_{1}}x_{im_{3}}s_{m_{3}k_{2}}x_{jm_{4}}s_{m_{4}k_{2}} \right)=E\left( x_{im_{1}}x_{im_{2}}x_{jm_{1}}x_{jm_{2}}s_{m_{1}k_{1}}s_{m_{2}k_{1}}s_{m_{1}k_{2}}s_{m_{2}k_{2}} \right)$$

**IV-1** $k_{1}=k_{2}$:

$$E\left( x_{im_{1}}x_{im_{2}}x_{jm_{1}}x_{jm_{2}}s_{m_{1}k_{1}}^{2}s_{m_{2}k_{1}}^{2} \right)=E\left( x_{im_{1}}x_{jm_{1}} \right)E\left( x_{im_{2}}x_{jm_{2}} \right)E\left( s_{m_{1}k_{1}}^{2} \right)E\left( s_{m_{2}k_{1}}^{2} \right)=\frac{\theta^{2}}{M^{2}}$$

|  | $\boldsymbol{M}$ **(counts)** |  | $\boldsymbol{K}$ **(counts)** | **Expectation** |
| --- | --- | --- | --- | --- |
| **(I)** | $m_{1}=m_{2}=m_{3}=m_{4} (M)$ | **(I-1)** | $k_{1}=k_{2} (K)$ | $\frac{3+6\theta^{2}}{M^{2}}$ |
|  |  | **(I-2)** | $k_{1}\neq k_{2} (K(K-1))$ | $\frac{1+2\theta^{2}}{M^{2}}$ |
| **(II)** | $m_{1}=m_{2}, m_{3}=m_{4} (M(M-1))$ | **(II-1)** | $k_{1}=k_{2} (K)$ | $\frac{\theta^{2}}{M^{2}}$ |
|  |  | **(II-2)** | $k_{1}\neq k_{2} (K(K-1))$ | $\frac{\theta^{2}}{M^{2}}$ |
| **(III)** | $m_{1}=m_{3}, m_{2}=m_{4} (M(M-1))$ | **(III-1)** | $k_{1}=k_{2} (K)$ | $\frac{1}{M^{2}}$ |
|  |  | **(III-2)** | $k_{1}\neq k_{2} (K(K-1))$ | $0$ |
| **(IV)** | $m_{1}=m_{4}, m_{2}=m_{4} (M(M-1))$ | **(IV-1)** | $k_{1}=k_{2} (K)$ | $\frac{\theta^{2}}{M^{2}}$ |
|  |  | **(IV-2)** | $k_{1}\neq k_{2} (K(K-1))$ | $0$ |

$$E\left( \hat{g}_{ij}^{2} \right)=\frac{1}{K^{2}M^{2}}\left[ 3MK+MK\left( K-1 \right)+MK\left( M-1 \right) \right]+\frac{\theta^{2}}{K^{2}M^{2}}\left[ 6MK+2MK\left( K-1 \right)+M\left( M-1 \right)K^{2}+MK\left( M-1 \right) \right]$$

$$=\frac{1}{K^{2}M^{2}}\left[ MK+MK^{2}+M^{2}K \right]+\frac{\theta^{2}}{K^{2}M^{2}}\left[ 3MK+MK^{2}+M^{2}K+M^{2}K^{2} \right]$$

Integrating them together gives,

$$var\left( \hat{g}_{ij}^{2} \right)=E\left( \hat{g}_{ij}^{2} \right)-E^{2}\left( \hat{g}_{ij} \right)$$

$$=\frac{1}{K^{2}M^{2}}\left[ MK+MK^{2}+M^{2}K \right]+\frac{\theta^{2}}{K^{2}M^{2}}\left[ 3MK+MK^{2}+M^{2}K \right]$$

$$=\frac{1+M+K+\theta^{2}\left( 3+M+K \right)}{MK}$$

$$\simeq\frac{M+K+\theta^{2}\left( M+K \right)}{MK}$$

$$=\frac{1+\theta^{2}}{M}+\frac{1+\theta^{2}}{K}$$

When considering LD between SNPs,

|  | $\boldsymbol{M}$ **(counts)** |  | $\boldsymbol{K}$ **(counts)** | **Expectation** |
| --- | --- | --- | --- | --- |
| **(I)** | $m_{1}=m_{2}=m_{3}=m_{4} (M)$ | **(I-1)** | $k_{1}=k_{2} (K)$ | $\frac{3+6\theta^{2}}{M^{2}}$ |
|  |  | **(I-2)** | $k_{1}\neq k_{2} (K(K-1))$ | $\frac{1+2\theta^{2}}{M^{2}}$ |
| **(II)** | $m_{1}=m_{2}, m_{3}=m_{4} (M(M-1))$ | **(II-1)** | $k_{1}=k_{2} (K)$ | $\frac{\theta^{2}+\rho^{2}+\theta^{2}\rho^{2}}{M^{2}}$ |
|  |  | **(II-2)** | $k_{1}\neq k_{2} (K(K-1))$ | $\frac{\theta^{2}+\rho^{2}+\theta^{2}\rho^{2}}{M^{2}}$ |
| **(III)** | $m_{1}=m_{3}, m_{2}=m_{4} (M(M-1))$ | **(III-1)** | $k_{1}=k_{2} (K)$ | $\frac{1+2\theta^{2}\rho^{2}}{M^{2}}$ |
|  |  | **(III-2)** | $k_{1}\neq k_{2} (K(K-1))$ | $0$ |
| **(IV)** | $m_{1}=m_{4}, m_{2}=m_{4} (M(M-1))$ | **(IV-1)** | $k_{1}=k_{2} (K)$ | $\frac{\theta^{2}+\rho^{2}+\theta^{2}\rho^{2}}{M^{2}}$ |
|  |  | **(IV-2)** | $k_{1}\neq k_{2} (K(K-1))$ | $0$ |

$$var\left( \hat{g}_{ij}^{2} \right)=E\left( \hat{g}_{ij}^{2} \right)-E^{2}\left( \hat{g}_{ij} \right)$$

$$=\frac{1+M+K+\theta^{2}\left( 3+M+K \right)}{MK}+\frac{1}{K^{2}M^{2}}\left[ K^{2}\sum_{m}^{M} \sum_{m^{'}}^{M} \left( \theta^{2}+\rho^{2}+\theta^{2}\rho^{2} \right)+K\sum_{m}^{M} \sum_{m^{'}}^{M} \left( 1+2\theta^{2}\rho^{2} \right)+K\sum_{m}^{M} \sum_{m^{'}}^{M} \left( \theta^{2}+\rho^{2}+\theta^{2}\rho^{2} \right) \right]$$

$$=\frac{1+M+K+\theta^{2}\left( 3+M+K \right)}{MK}+\frac{1}{K^{2}M^{2}}\left[ K\left( K+1 \right)\left( \frac{M^{2}}{M_{e}}-M \right)+K\left( K+3 \right)\left( \frac{M^{2}}{M_{e}}-M \right)\theta^{2} \right]$$

$$=\frac{1+M+K+\theta^{2}\left( 3+M+K \right)}{MK}+\frac{\left( M-M_{e} \right)\left[ K+1+\left( K+3 \right)\theta^{2} \right]}{KMM_{e}}$$

$$=\frac{1+M_{e}+K+\theta^{2}\left( 3+M_{e}+K \right)}{M_{e}K}$$

$$\simeq\frac{1+\theta^{2}}{M_{e}}+\frac{1+\theta^{2}}{K}$$

#### Note 5 The variance of encG-reg

For a simple regression model, $y=bx+e$. Least square estimator of $b$ is $\hat{b}=\frac{cov\left( x,y \right)}{var\left( x \right)}$. The variance of regression coefficient is $\sigma_{b}^{2}\simeq\frac{\sigma_{y}^{2}\left( 1-\rho^{2} \right)}{n\sigma_{x}^{2}}$, where $n$ is the number of observation and $\rho^{2}=\frac{\sigma^{2}\left( x,y \right)}{\sigma_{x}^{2}\sigma_{y}^{2}}$. In encG-reg, if we assign $y=\hat{x}_{ik}$ and $x=\hat{x}_{jk}$, each of which is a $K\times$1 vector, the estimator $\hat{b}_{ij}=\frac{cov\left( \hat{x}_{ik},\hat{x}_{jk} \right)}{var\left( \hat{x}_{jk} \right)}=cov\left( \hat{x}_{ik},\hat{x}_{jk} \right)$. Then, the variance of regression coefficient is,

$$\sigma_{b}^{2}\simeq\frac{1-\rho^{2}}{K}=\frac{1-\sigma^{2}\left( \hat{x}_{ik},\hat{x}_{jk} \right)}{K}\simeq\frac{1-\theta^{2}}{K}$$

$$\hat{b}_{ij}=cov\left( \hat{x}_{ik},\hat{x}_{jk} \right)=cov\left( \sum_{m}^{M} x_{im}s_{mk},\sum_{m}^{M} x_{jm}s_{mk} \right)=\sum_{m}^{M} cov\left( x_{im}s_{mk},x_{jm}s_{mk} \right)$$

$$E\left( \hat{b}_{ij} | \hat{\theta} \right)=E\left( \hat{b}_{ij} \right|x_{im},x_{jm})=\sum_{m}^{M} x_{im}x_{jm}var(s_{mk})=\frac{1}{M}\sum_{m}^{M} x_{im}x_{jm}=g_{ij}$$

$$var\left( \hat{b}_{ij} \right)=E\left[ var\left( \hat{b}_{ij} | \hat{\theta} \right) \right]+var\left[ E\left( \hat{b}_{ij} | \hat{\theta} \right) \right]=E\left( \frac{1-\hat{\theta}^{2}}{K} | \hat{\theta} \right)+var\left( g_{ij} \right)\simeq\frac{1-\hat{\theta}^{2}}{K}+\frac{1-\theta^{2}}{M}$$

#### Note 6 Details about power calculation in Eq 4

For hypothesis on a statistic $T$, which follows $T\sim N\left( \mu_{0}, \sigma_{0}^{2} \right)$ for $H_{0}$ and follows $T\sim N\left( \mu_{1}, \sigma_{1}^{2} \right)$ for $H_{1}$. We assume that $\sigma_{0}^{2}=f_{0}^{2}/n$ and $\sigma_{1}^{2}=f_{1}^{2}/n$, where $n$ is the number of sample size.

For two-tailed hypothesis that $H_{0}: \mu=\mu_{0}$, and $H_{1}: \mu\neq\mu_{0}$, the minimal number of $n$ is,

$$n>\left( \frac{z_{1-\beta}f_{1}+z_{1-\alpha/2}f_{0}}{\mu_{1}-\mu_{0}} \right)^{2}$$

In this case, $H_{0}: \mu_{0}=\theta_{r}=0$, and $H_{1}: \mu_{1}=\theta_{r}$. Given $\sigma_{0}^{2}=\frac{1}{k}+\frac{1}{m}=\frac{1}{\frac{km}{k+m}}$ and $\sigma_{1}^{2}=\frac{1-\theta_{r}^{2}}{k}+\frac{1-\theta_{r}^{2}}{m}=\frac{1-\theta_{r}^{2}}{\frac{km}{k+m}}$, the minimal number of $\frac{km}{k+m}$ is,

$$\frac{km}{k+m}>\left( \frac{z_{1-\beta}\sqrt{1-\theta_{r}^{2}}+z_{1-\alpha/2}}{\theta_{r}} \right)^{2}$$

$$\frac{1}{k}+\frac{1}{m}<\frac{1}{\left( \frac{z_{1-\beta}\sqrt{1-\theta_{r}^{2}}+z_{1-\alpha/2}}{\theta_{r}} \right)^{2}}$$

$$k>\frac{1}{\left( \frac{\theta_{r}}{z_{1-\beta}\sqrt{1-\theta_{r}^{2}}+z_{1-\alpha/2}} \right)^{2}-\frac{1}{m}}$$

In addition, $m$ can be replaced with $m_{e}$ if LD is taken into account.

**Note 7 Selection of ethnicity-insensitive SNPs.**

Typically, one of the focuses in population genetics is identifying genetic variants associated with population structure, which are referred to as ethnicity-sensitive SNPs. In contrast, encG-reg is interested in SNPs with similarly distributed allele frequencies across different populations worldwide, which we call ethnicity-insensitive SNPs. We selected ethnicity-insensitive SNPs based three reference populations from 1000 Genome Project, referred to as 1KG-CHN (103 CHB and 105 CHS samples), 1KG-EUR (99 CEU and 107 TSI samples), and 1KG-AFR (107 YRI samples), respectively. YRI represents African samples, CHB and CHS represent Han Chinese in Beijing and Southern Han Chinese, CEU represents Utah residents with Northern and Western European ancestry, and TSI represents Tuscani in Italy.

We first performed SNP quality control on 1KG-CHN, 1KG-EUR, and 1KG-AFR, using the following inclusion criteria: (1) minor allele frequency (MAF) > 0.01; (2) Hardy-Weinberg equilibrium (HWE) test p-value > 1e-7; and (3) locus missingness < 0.05. Next, we conducted association studies on two reference populations at a time, selecting SNPs with a p-value greater than 0.05 (considered insignificant). Once insignificant SNPs were identified between each pair of reference populations, we took the intersection between those SNPs to establish the ethnicity-insensitive SNP pool, which contained a total of 299,835 SNPs. These SNPs exhibit consistent allele frequency across different ethnic backgrounds.

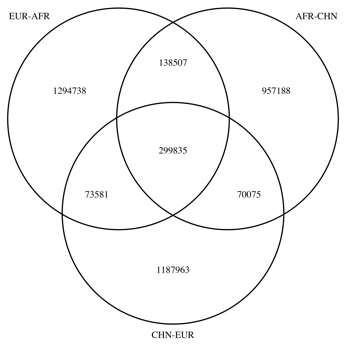

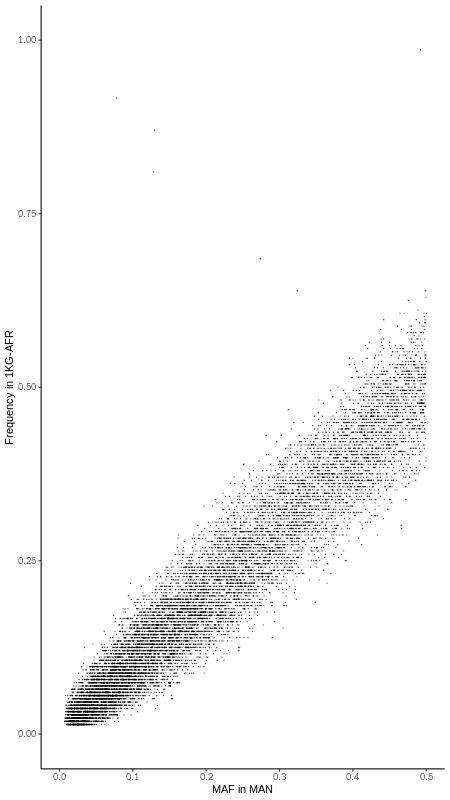

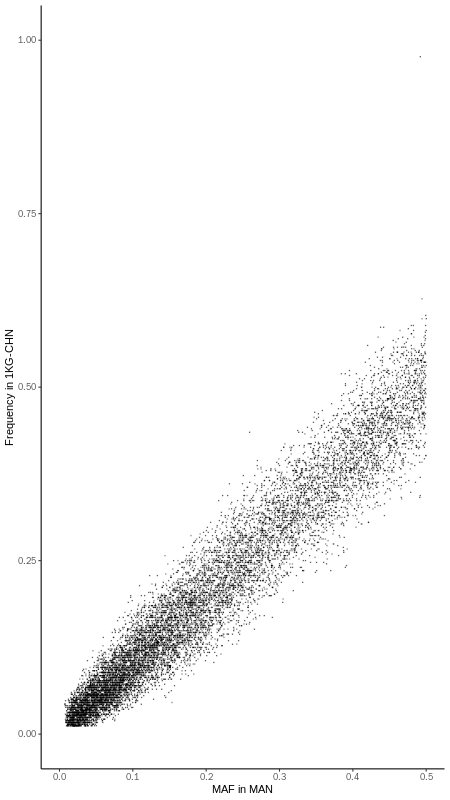

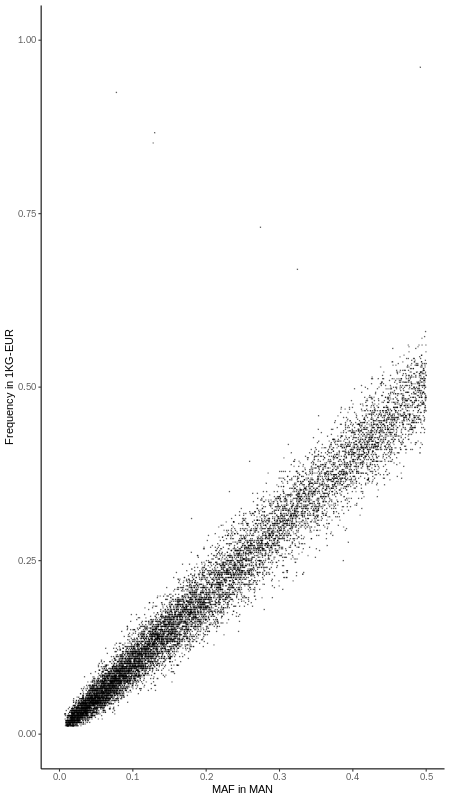

**Note 8 fPCA and fStructure**

fPCA is a PCA calculation method based on summary data at the population level, rather than individual-level data. It uses the allele frequency of common markers across all cohorts to construct a frequency matrix $\mathbf{P}=\left\{ p_{ij} \right\}_{\mathcal{C\times M}}$ for PCA calculation. Here, $\mathcal{C}$ represents the number of cohorts and $\mathcal{M}$ represents the number of common markers, while $p_{ij}$ denotes the allele frequency for the $j$th marker in the $i$th cohort. By analyzing the differences in marker frequencies, fPCA can effectively capture the population structure in PC1 and PC2.

fStructure explores the genetic composition of the target cohorts by comparing with reference populations using $F_{st}$. Again, $F_{st}$ is calculated based on allele frequency of common markers among all cohorts and reference population. Suppose there are $\mathcal{C}$ cohorts and three reference population. The average $F_{st}$ using common markers ($F_{st}^{ref1}$, $F_{st}^{ref2}$, and $F_{st}^{ref3}$) between $i$th cohort and the given reference population are calculated. Finally, a bar plot is employed to show the ratio of $1/F_{st}^{ref1}$, $1/F_{st}^{ref2}$, and $1/F_{st}^{ref3}$, providing a clear visualization of the genetic composition of the target cohort against the reference populations.
