## Supplementary material for "Searching across-cohort relatives in 54,092 GWAS samples via encrypted genotype regression": SuppNoteFigsTables: S2_Table.docx

### S2 Table. $\boldsymbol{m}_{\boldsymbol{e}}$ estimation in UK Biobank and Chinese cohorts.

| $\boldsymbol{m}$ |  | ${\hat{\boldsymbol{m}}}_{\boldsymbol{e}}$ | |
| --- | --- | --- | --- |
|  |  | **Manchester** | **Oxford** |
| 500 |  | 494 | 495 |
| 1,000 |  | 967 | 968 |
| 5,000 |  | 4,398 | 4,418 |
| 10,000 |  | 8,024 | 8,067 |

| $\boldsymbol{m}$ |  | ${\hat{\boldsymbol{m}}}_{\boldsymbol{e}}$ | | | |
| --- | --- | --- | --- | --- | --- |
|  |  | **1KG-CHN** | **UKB-CHN** | **CONVERGE** | **MESA** |
| 500 |  | 477 | 454 | 465 | 472 |
| 1,000 |  | 970 | 893 | 923 | 915 |
| 5,000 |  | 4,190 | 3,830 | 4,116 | 3,798 |
| 10,000 |  | 7,355 | 6,648 | 7,323 | 6,991 |

**Table Notes**: For all filtered SNPs, different number of SNPs ($m$=500, 1,000, 5,000, 10,000) are used to test the size of $m_{e}$ in homogenous cohorts, taking cohorts in UKB (Manchester and Oxford) and in China (1KG-CHN, UKB-CHN, CONVERGE and MESA) as employed in this study.
