## Supplementary material for "Searching across-cohort relatives in 54,092 GWAS samples via encrypted genotype regression": SuppNoteFigsTables: S3_Table.docx

### S3 Table. Sample size for 19 cohorts in UKB.

| **Cohort** | **Sample size** |
| --- | --- |
| Barts | 12,393 |
| Birmingham | 24,510 |
| Bristol | 42,290 |
| Bury | 27,957 |
| Cardiff | 17,666 |
| Croydon | 26,412 |
| Edinburgh | 16,674 |
| Glasgow | 18,287 |
| Hounslow | 27,664 |
| Leeds | 43,421 |
| Liverpool | 32,031 |
| Manchester | 13,799 |
| Middlesbrough | 20,757 |
| Newcastle | 36,224 |
| Nottingham | 33,386 |
| Oxford | 13,911 |
| Reading | 29,114 |
| Sheffield | 29,531 |
| Stoke | 19,131 |
| **Total number of samples** | **485,158** |
| **Total number of inter-cohort paired samples** | **110,713,926,381** |
