## Supplementary material for "Searching across-cohort relatives in 54,092 GWAS samples via encrypted genotype regression": SuppNoteFigsTables: S5_Table.docx

### S5 Table. The number of inferred relatedness at exhaustive and parsimony designs.

| **Design^a^** | **Method** | **Number of inferred relatedness** | | | **Total** |
| --- | --- | --- | --- | --- | --- |
|  |  | **Identical** | **1st-degree** | **2nd-degree** |  |
| Exhaustive design | KING | 38 | 7,965 | 6,632 | 14,635 |
|  | encG-reg | 38 | 7,913 | 7,022 | 14,973 |
| Parsimony design | encG-reg | 38 | 7,903 | 5,435 | 13,376 |

^a^Exhaustive design denotes the use of the intersected SNPs between each pair of cohorts and parsimony design denotes the use of intersected SNPs of all participated cohorts. When performing encG-reg at 19 cohorts in UKB, experiment-wise Bonferroni correction is based on the number of paired samples between each two cohorts ($n_{i}\times n_{j}$) for exhaustive design and based on total number of paired samples among all cohorts ($\mathcal{N=}\sum_{i<j}^{\mathcal{C}} n_{i}n_{j}$, where there are $\mathcal{C}$ cohorts and $n_{i}$ is the sample size of cohort $i$ for parsimony design.
