## Supplementary material for "Searching across-cohort relatives in 54,092 GWAS samples via encrypted genotype regression": SuppNoteFigsTables: S6_Table.docx

### S6 Table. Summary of cross-cohort quality control.

| **Cohort ID** | **Input** | **No palindrome** | **Intersect with CONVERGE** | **After removing flipped SNPs** |
| --- | --- | --- | --- | --- |
| CONVERGE | 5,215,820 | 5,215,820 |  |  |
| SBWCH | 1,237,941 | 1,237,941 | 1,201,528 | 1,201,528 |
| CAS1 | 288,684 | 288,684 | 256,950 | 256,944 |
| CAS2 | 288,539 | 288,539 | 256,938 | 256,932 |
| Fudan | 311,384 | 311,384 | 310,596 | 310,596 |
| WBBC | 319,930 | 319,930 | 285,956 | 285,897 |
