## Supplementary figures and images for "Searching across-cohort relatives in 54,092 GWAS samples via encrypted genotype regression"

### S1_Fig.pdf

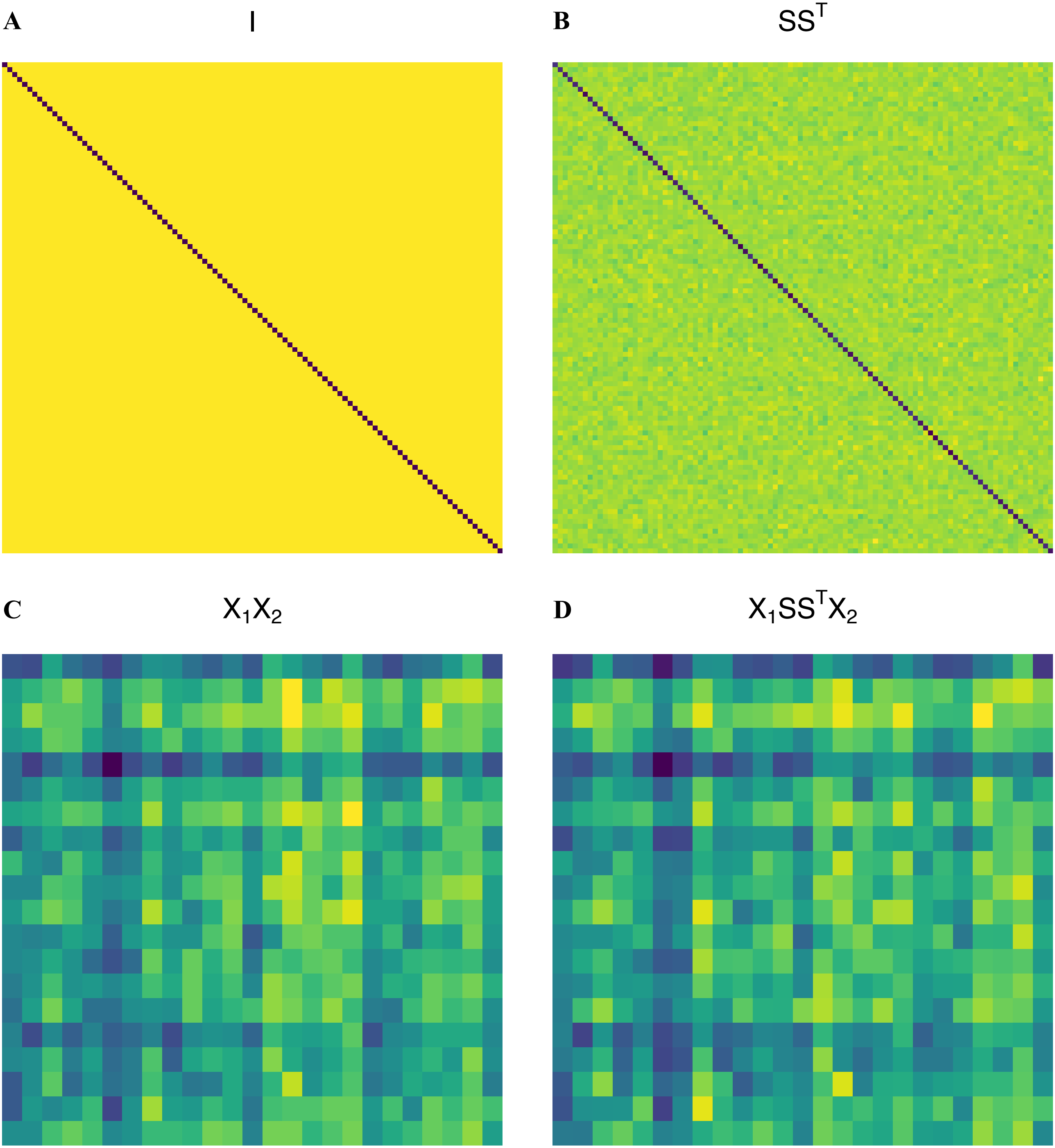

### S2_Fig.pdf

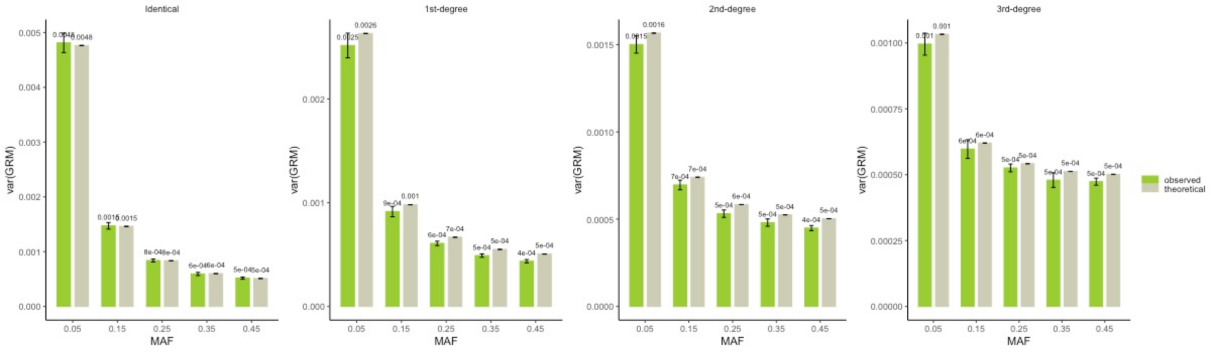

### S3_Fig.pdf

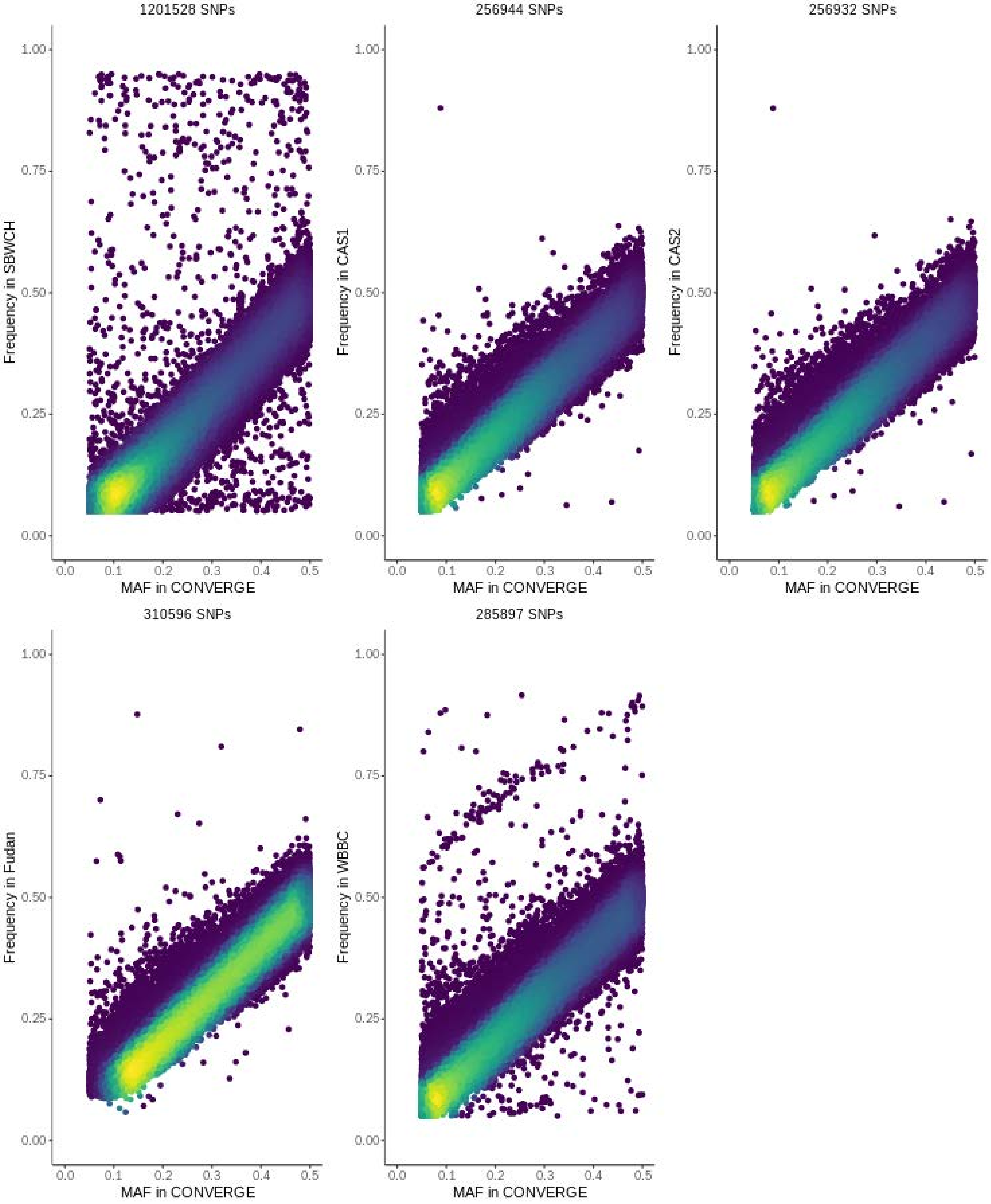

### S4_Fig.pdf

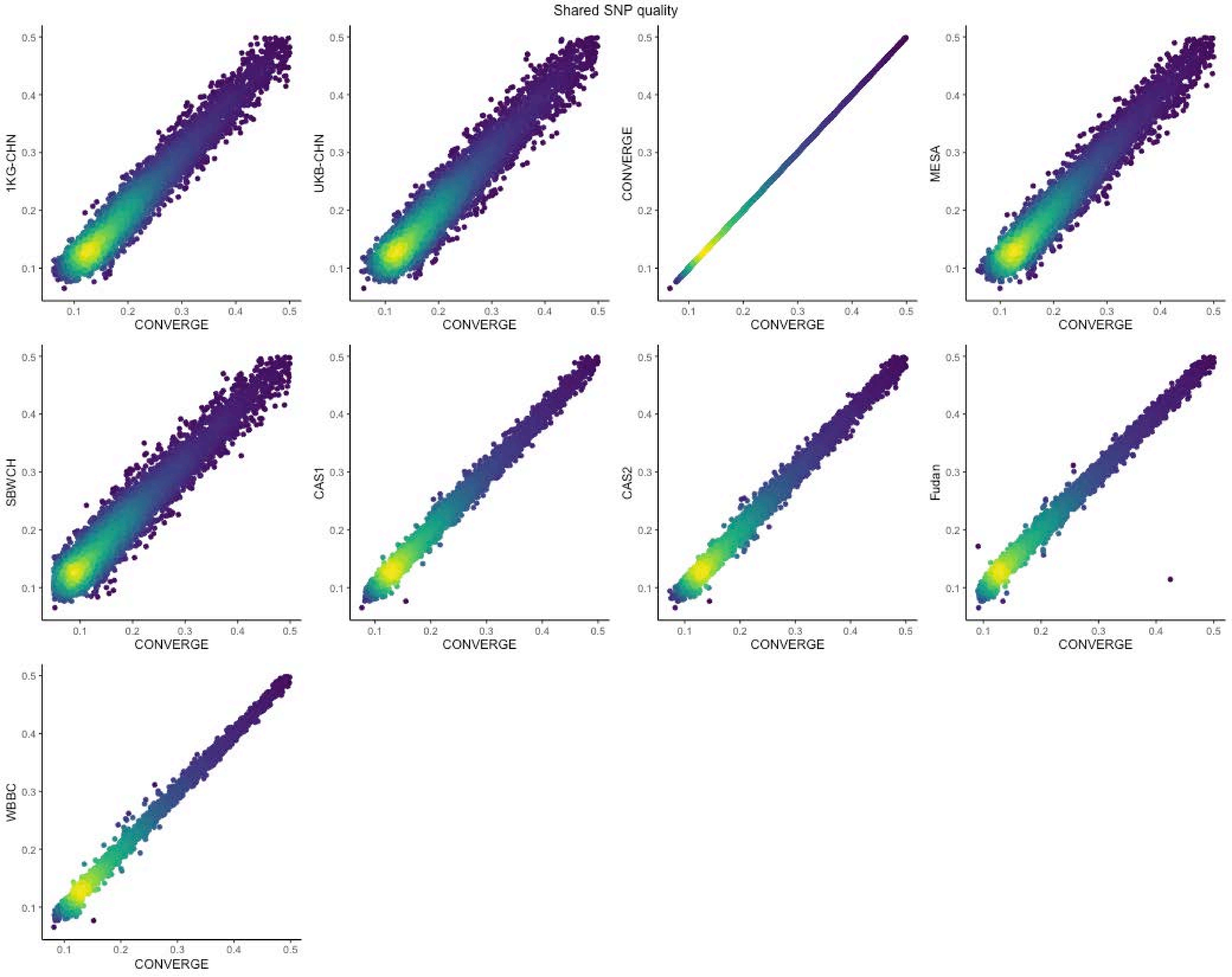

### S5_Fig.pdf

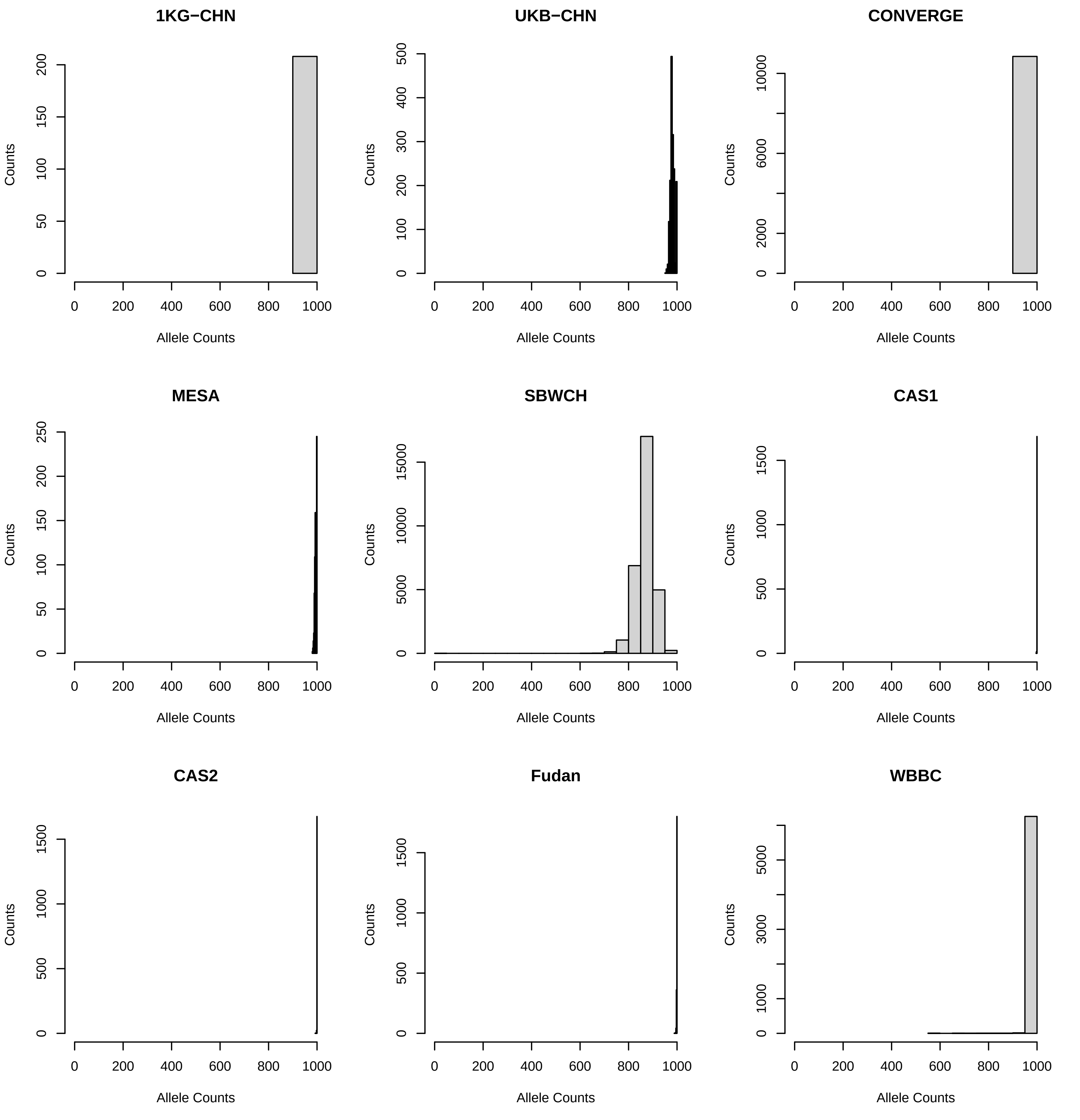

### S6_Fig.pdf

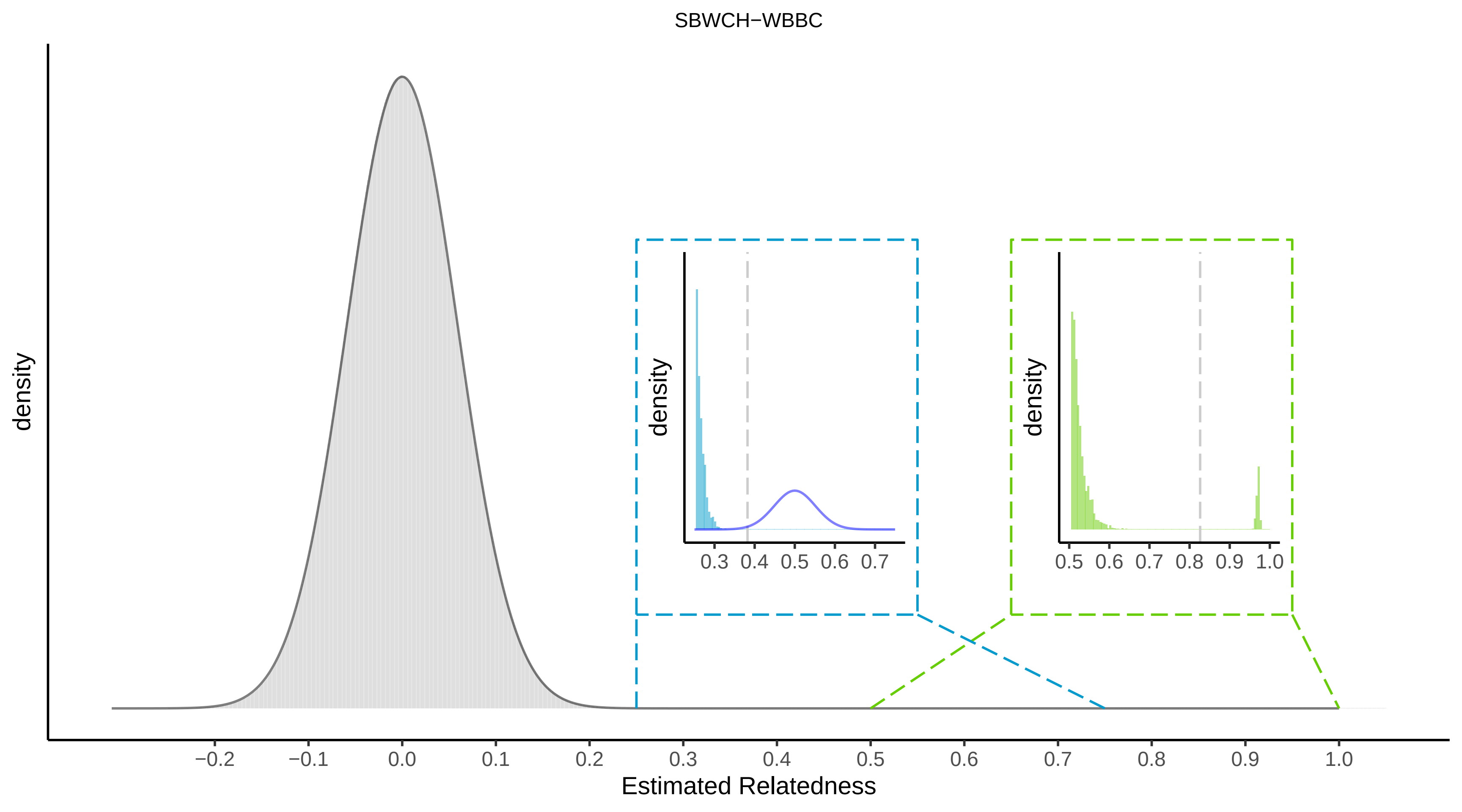
